# Usefulness Criterion and post-selection Parental Contributions in Multi-parental Crosses: Application to Polygenic Trait Introgression

**DOI:** 10.1101/484287

**Authors:** Antoine Allier, Laurence Moreau, Alain Charcosset, Simon Teyssèdre, Christina Lehermeier

**Author notes:** Corresponding authors Christina Lehermeier, RAGT 2n, Genetics & Analytics Unit, 12510 Druelle, France, Simon Teyssèdre, RAGT 2n, Genetics & Analytics Unit, 12510 Druelle, France.

## Abstract

Predicting the usefulness of crosses in terms of expected genetic gain and genetic diversity is of interest to secure performance in the progeny and to maintain long-term genetic gain in plant breeding. A wide range of crossing schemes are possible including large biparental crosses, backcrosses, four-way crosses, and synthetic populations. *In silico* progeny simulations together with genome-based prediction of quantitative traits can be used to guide mating decisions. However, the large number of multi-parental combinations can hinder the use of simulations in practice. Analytical solutions have been proposed recently to predict the distribution of a quantitative trait in the progeny of biparental crosses using information of recombination frequency and linkage disequilibrium between loci. Here, we extend this approach to obtain the progeny distribution of more complex crosses including two to four parents. Considering agronomic traits and parental genome contribution as jointly multivariate normally distributed traits, the usefulness criterion parental contribution (UCPC) enables to (i) evaluate the expected genetic gain for agronomic traits, and at the same time (ii) evaluate parental genome contributions to the selected fraction of progeny. We validate and illustrate UCPC in the context of multiple allele introgression from a donor into one or several elite recipients in maize (*Zea mays* L.). Recommendations regarding the interest of two-way, three-way, and backcrosses were derived depending on the donor performance. We believe that the computationally efficient UCPC approach can be useful for mate selection and allocation in many plant and animal breeding contexts.

## INTRODUCTION

Allocation of resources is a key factor of success in plant and animal breeding. At each selection cycle, breeders are facing the choice of crosses to generate the genetic variation on which selection will act at the next generation. In case of limited genetic variation for targeted traits, the introduction of favorable alleles from donors to elite material is necessary to ensure long term genetic gain. Several approaches have been proposed to introgress superior quantitative trait locus (QTL) alleles from a donor into a recipient. In case of a single desirable allele, it can be accomplished using molecular assisted introgression (Visscher *et al.* 1996; Frisch *et al.* 1999). In case of multiple desirable alleles, gene pyramiding strategies have been proposed (Hospital and Charcosset 1997; Charmet *et al.* 1999; Servin *et al.* 2004). More recently, Han et al. (2017) proposed the predicted cross value (PCV) to select at each generation crosses that maximize the likelihood of pyramiding desirable alleles in their progeny. For quantitative traits implying numerous QTL with small individual effects, genomic selection has been proposed to fasten the introgression of exotic alleles into elite germplasm (Bernardo 2009) and to harness polygenic variation from genetic resources (Gorjanc *et al.* 2016) using two-way crosses or backcrosses. However, plant breeders are not only considering biparental crosses such as two-way crosses or backcrosses but also multi-parental crosses including three-way crosses, four-way crosses or synthetic populations (Gallais 1990; Schopp *et al.* 2017). Crosses implying several parental lines are highly interesting for breeders to exploit at best the genetic diversity underlying one or several traits. Beyond fastening the introgression of genetic resources into elite germplasm, genomic selection could be used to predict the interest of a multi-parental cross involving one or several donors and recipients. Among possible crosses, the identification of those that secure the performance in progeny and maximize the genome contribution of donors to the selected progeny is essential for increasing or maintaining genetic gain and diversity of an elite population.

The interest of a cross for a given quantitative trait can be defined using the usefulness criterion (Schnell and Utz 1975) that is determined by its expected genetic mean (μ) and genetic gain (*ihσ*): *UC* = μ + *i h σ*, where *σ* is the progeny genetic standard deviation. The selection intensity (*i*) depends on the selection pressure and the selection accuracy (*h*) can be assumed to be one when selecting on genotypic effects (Zhong and Jannink 2007). While μ can be easily predicted for different crossing schemes by the weighted average of parental values, the difficulty to have a good prediction of progeny variance (*σ*^2^) hindered the use of UC in favor of simpler criteria (for a recent review on different criteria, see Mohammadi *et al.* 2015). Bernardo *et al.* (2006) suggested to predict the progeny variance of a given population using genotypic data of its progenitors and quantitative trait loci (QTL) effect estimates, assuming unlinked QTLs. Zhong and Jannink (2007) extended this concept to linked loci. With the availability of high-density genotyping, it has been proposed to predict the progeny variance using *in silico* simulations of progeny and genome-wide marker effects (Bernardo 2014; Lian *et al.* 2015; Mohammadi *et al.* 2015). However, the geometrically increasing number of cross combinations possible for *n* parents makes the testing of all crosses computationally intensive. For instance, with only *n* = 50 potential parents, a total of 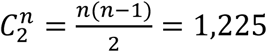 genetically different two-way crosses can be formed. This number increases by a factor of *n* when crossing all the possible two-way crosses to the *n* different parents, so that 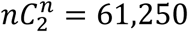 three-way crosses and backcrosses are possible. Recently, Lehermeier *et al.* (2017b) derived algebraic formulas to predict for a single trait the genetic variance of doubled haploid (DH) or recombinant inbred line (RIL) progeny derived from two-way crosses, using information of recombination frequency and linkage disequilibrium in parental lines. These algebraic formulas have not been extended so far to multi-parental crosses, hindering the prediction of the interest of such crosses.

While the expected genetic gain (UC) is a meaningful measure of the interest of a cross for breeding, it does not account for the parental genome contributions to the selected fraction of progeny that determine the genetic diversity in the next generation. Parental genome contribution to unselected progeny has been studied for several years and is of specific interest in breeding for donor introduction and to manage long term genetic gain and inbreeding rate (Hill 1993; Bijma 2000; Woolliams *et al.* 2015). Hill (1993) derived the variance of the non-recurrent parent genome contribution to heterozygous backcross individuals in cattle. Wang and Bernardo (2000) formulated the variance of parental genome contribution to F2 and backcross plant progeny considering a finite number of loci. Frisch and Melchinger (2007) extended this approach to a continuous integration over loci and showed that a normal distribution approximated well parental genome contribution obtained from computer simulations. Also empirical data on pairs of human full-sibs confirmed that parental genome contributions, i.e. additive relationship, can be considered as normally distributed around the expected value of 0.5 (Visscher *et al.* 2006; Visscher 2009). All these studies considered the parental genome contribution distribution in unselected progeny. However, to control parental contribution during polygenic traits introgression, it is of interest to predict parental genome contribution after selection for quantitative traits.

In this study, we develop a multivariate approach called usefulness criterion parental contribution (UCPC) to evaluate the interest of a multi-parental cross implying a donor line and one or several elite recipients based on the expected genetic gain (UC) and the diversity (parental contributions, PC) in the selected progeny. We extend here the rational given by Lehermeier *et al.* (2017b) for two important aspects. We address the prediction of progeny variance for multi-parental crosses implying two to four parents and we consider the parental contribution as an additional quantitative trait. The originality of this approach is that it uses derivations of the prediction of progeny variance in multi-parental crosses implying up to four parents to jointly predict (i) the performance of the next generation using the usefulness criterion and (ii) the parental contributions to the selected fraction of progeny, which to our knowledge has not been investigated so far. We illustrate the use of UCPC in the context of external genetic resources introgression into elite material considering the specific case of a unique donor that is crossed to one or several elite recipients. We address the type of multi-parental cross that should be preferred among two-way crosses, three-way crosses or backcrosses in order to maximize genetic gain while introgressing donor alleles in the elite population within one selection cycle.

## MATERIALS AND METHODS

### Application example: breeding context

We assumed a generic plant breeding population of fully homozygote inbred lines genotyped for biallelic single nucleotide polymorphism (SNP) markers with known positions. We considered a quantitative agronomic trait (e.g. grain yield) implying *p* QTLs with known additive effects and with positions sampled among the SNP marker positions. Further, we considered that the breeding population is an elite population that should be enriched with alleles from a donor. We assumed a donor line (D) has been identified and should be crossed with lines from the elite population (e.g. E1 and E2) in order to obtain high-performing progeny that combine donor favorable alleles in a performing elite background. This donor line can vary in its performance level and its diversity relative to the elite population.

In this context, we aimed at evaluating the interest of two-way crosses (i.e. *D* × *E*_1_ and *D* × *E*_2_), backcrosses (i.e. (*D* × *E*_1_) × *E*_1_ and (*D* × *E*_2_) × *E*_2_) or three-way crosses (i.e. (*D* × *E*_1_) × *E*_2_ and (*D* × *E*_2_) × *E*_1_) based on (i) the mean performance of the selected progeny and (ii) the average genome contribution of the donor to the selected progeny. Considering different donor characteristics, i.e. originality and performance level, we compared the interest of the multi-parental crosses listed above in order to derive guidelines for the use of the donor *D*. As a benchmark, we also evaluated the interest of different elite multi-parental crosses.

### Usefulness Criterion Parental Contribution

In order to predict the progeny distribution of a given cross in terms of expected genetic gain and genetic diversity, we considered the agronomic trait and the parental genome contribution as jointly multivariate normally distributed traits. This enabled us to (i) evaluate the genetic gain of the selected progeny for the agronomic trait, and to (ii) evaluate the contribution of each parental line to this selected progeny. An illustration of the concept of UCPC is given in Figure 1. In the following sections we present in more detail the theory underlying UCPC in the general case of a four way cross.

**Figure 1.**
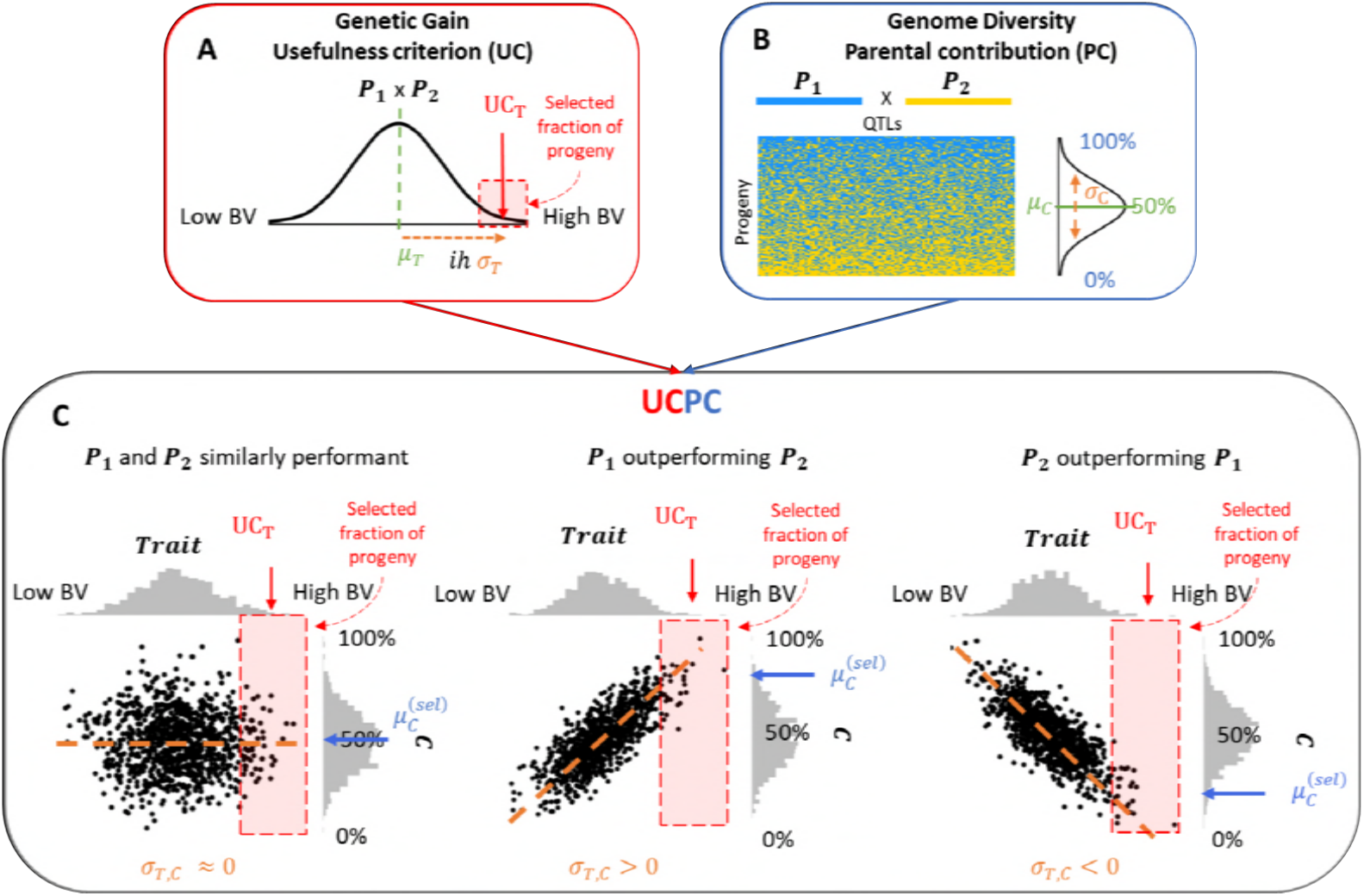
Illustration of Usefulness Criterion Parental Contribution (UCPC) for a two-way cross between *P*_1_ and *P*_2_. UCPC combines (A) the concept of usefulness criterion for an agronomic trait normally distributed (*N*(μ_*T*_, *σ*_*T*_)) and (B) *P*_1_ genome contribution considered as a normally distributed quantitative trait (*N*(μ_*C*_, *σ*_*C*_)) in a multivariate approach (C). UCPC enables to predict the expected progeny performance for the trait (*UC*_*T*_) and *P*_1_ genome contribution to the selected fraction on progeny 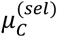 that depends on the covariance *σ* mainly driven by the difference between *P*_1_ and *P*_2_ performances.

### Multi-parental crosses and genetic model

To cover diverse types of crosses, we consider a general multi-parental cross implying four fully homozygous parents (*P*_1_, *P*_2_, *P*_3_ and *P*_4_, Figure 2). Note that for this general presentation of the theory, parents can be lines from the elite population and/or considered as external donors. This four-way cross implies two initial crosses giving generations 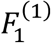 and 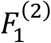, respectively (Figure 2). A second cross between 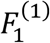 and 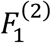 yields the generation *F*1′ standing for pseudo F1. Two-way crosses, three-way crosses and backcrosses can be seen as specific cases of four-way crosses depending on the number of parents considered as visualized in Figure 2.

**Figure 2.**
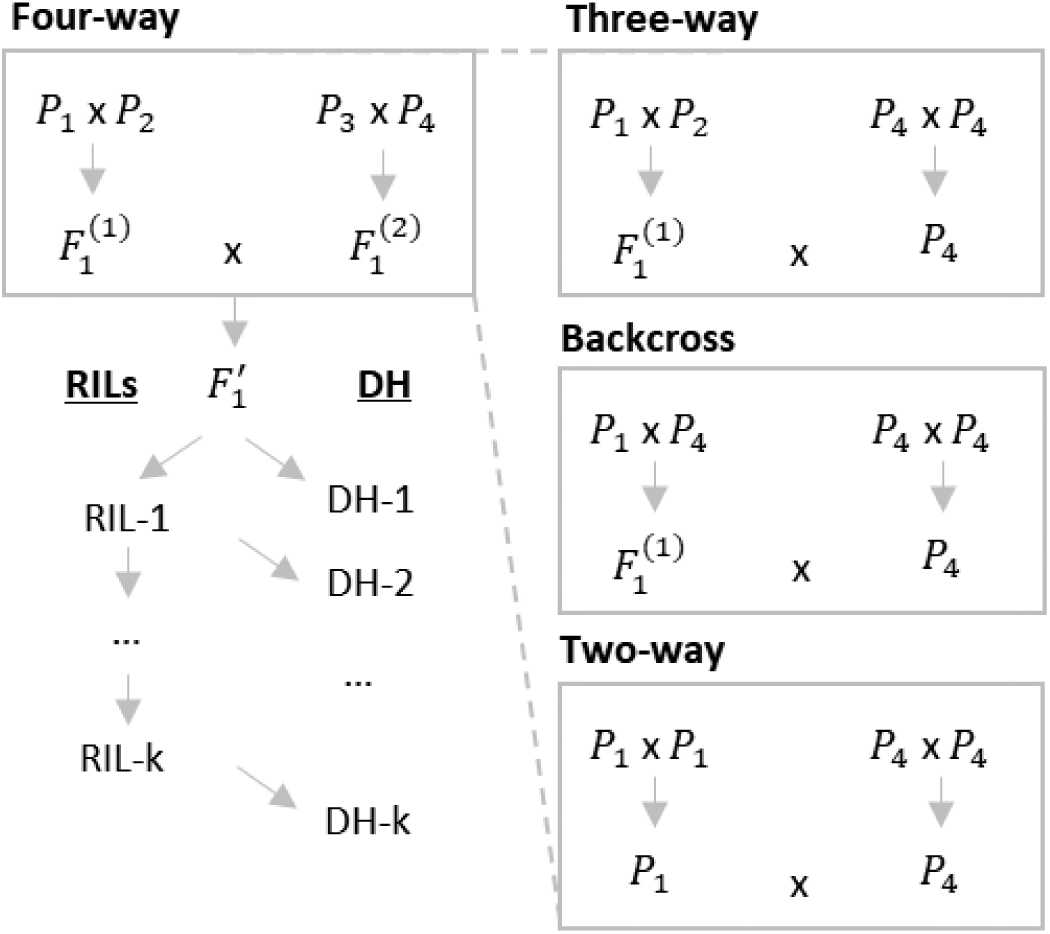
Illustration of four-way crosses (left) and derived crossing schemes (right). In the general case of four-way crosses, nomenclature is defined for recombinant inbred lines (RILs) after k generations of selfing (RIL-k) from pseudo F1 generation (F1’) and doubled haploid lines (DH) derived from the RIL generation k-1 (DH-k, for k > 1). RIL-1 corresponds to the pseudo F2 generation and RIL ∞ = DH ∞.

Assuming known genotypes at *p* QTLs underlying the quantitative trait considered and biallelic markers at QTL positions, ***x***_*i*_ denotes the *p*-dimensional genotype vector of parent *i*, with the *j*^th^ element coded as 1 or −1 for the genotypes AA or aa at locus *j*. Assuming biallelic QTL effects, a classical way to define the parental genotypes matrix would be a (4 × *p*)-dimensional matrix (***x***_1_ ***x***_2_ ***x***_3_ ***x***_4_)^′^. Addressing parental specific effects and following the identical by descent (IBD) genome contribution of parents to progeny requires to consider parental specific alleles. Thus, we extend the definition of parental genotypes to a multi-allelic coding:

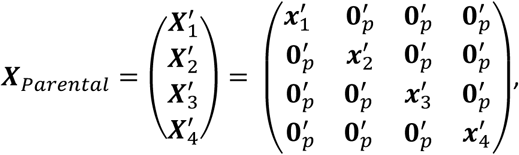

with ***X***_*Parental*_ a (4 × 4*p*) dimensional matrix defining the genotype of the four parents at the 4*p* parental alleles at QTLs, ***X***_*i*_ the 4*p*-dimensional vector defining the genotype of parent *i* and **0**_*p*_ a *p*-dimensional vector of zeros.

We first concentrate on doubled haploid (DH) lines derived from the *F1′* generation (DH-1), and then extend our work to DH lines generated after more selfing generations from the *F1′* and to recombinant inbred lines (RILs) at different selfing generations, i.e. partially heterozygous progeny. Absence of selection is assumed while deriving the progeny from generation *F1′*. In case of DH-1, we denote the (*N* × 4*p*)-dimensional genotyping matrix of *N* progeny derived from a four-way cross (Figure 2) in a multi-allelic context as:

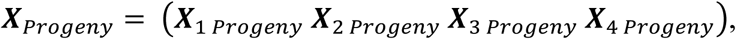

where for instance ***X***_1 *Progeny*_ is a (*N* × *p*)-dimensional matrix of progeny genotypes at QTLs coded −1 or 1 for alleles inherited from parent *P*_1_ and 0 otherwise.

The multi-parental coding enables to consider 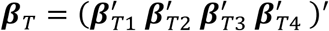 a 4*p*-dimensional vector of known parental specific additive effects for the agronomic trait. Thus, ***X***_*Progeny*_ ***β***_*T*_ is the vector of progeny breeding values of the agronomic trait. As we assumed additive effects, the breeding value equals the genetic value. Assuming no parental specific effects for the agronomic trait, as in the application example considered, ***β***_*T*_ reduces to 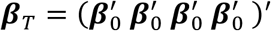, where ***β*_0_** is the vector of known QTL effects in the elite and donor populations. Furthermore, the multi-parental coding considered enables to define the effects to follow IBD parental contributions either genome-wide (namely *C*, ***β*_*C*_**) or considering only the favorable alleles (namely *C*(+), ***β*_*C*_**_(+)_). In this study, we focused on the first parent (*P*_1_) genome IBD contributions, but a generalization to every parent is straightforward. In the following, ***β***_*C*_ is a 4*p*-dimensional vector defined to follow *P*_1_ genome-wide contribution and ***β***_*C*(+)_ a 4*p*-dimensional vector defined to follow *P*_1_ genome contribution at favorable alleles. In the general case of four-way crosses 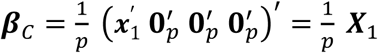 and ***β***_*C*(+)_ is identical to ***β***_*C*_ except that if *P*_1_ has the unfavorable allele at QTL *q* ∈ [1, *p*], the corresponding element of ***β***_*C*(+)_ is null. Thus, ***X***_*Progeny*_ ***β***_*C*_ represents the proportion of alleles in the progeny that are inherited from *P*_1_ independently of the allele effect and ***X***_*Progeny*_ ***β***_*C*(+)_ represents the proportion of alleles in the progeny that are inherited from *P*_1_ and favorable. In the specific case of two-way crosses (i.e. *P*_1_ = *P*_2_ and *P*_3_ = *P*_4_ so ***x***_1_ = ***x***_2_ and ***x***_4_ = ***x***_3_), *P*_1_ genome-wide contribution is defined by 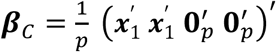.

### Prediction of progeny mean and progeny variance

In this section we consider a generic quantitative trait defined by the 4*p*-dimensional vector of parent specific additive effects 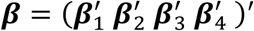. The vector ***β*** can be replaced by ***β***_*T*_, ***β***_*C*_ or ***β***_*C*(+)_ without loss of generality. In order to evaluate the performance of a four-way cross, we derive its expected progeny mean and variance. The expected progeny mean can be derived as the mean of all four parents’ breeding values:

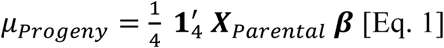

The progeny variance can be derived as:

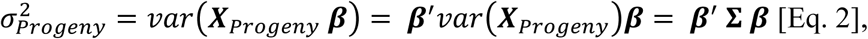

where **Σ** is the (4*p* × 4*p*)-dimensional covariance matrix between parental alleles at QTL in progeny. The diagonal elements **Σ**_*jj*_ (*j* ∈ [1,4*p*]) are equal to the variance of parental alleles in progeny. Note that off-diagonal elements **Σ**_*jl*_ (*j* ≠ *l* ∈ [1,4*p*]) correspond to the disequilibrium covariance between two parental alleles *j* and *l* at different QTLs (i.e. different physical positions) or at the same QTL. The linkage disequilibrium parameter in the progeny between parental alleles *D*_*jl*_ can be derived from the linkage disequilibrium parameter among the four parental lines and the recombination frequency between parental alleles in progeny (Table 1, see File S1 for derivation). In the specific case considered, i.e. doubled haploid lines derived from generation *F1′* (DH-1), this leads to the covariance entry:

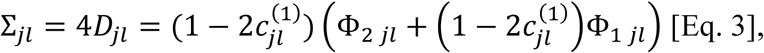

where 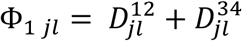 is the sum of the disequilibrium parameter between parental alleles *j* and *l* in pairs of parents implied in the first crosses and 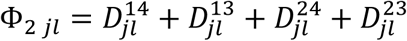 is the sum of disequilibrium parameter between parental alleles *j* and *l* in pairs of parents indirectly implied in the second cross. 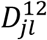 denotes the linkage disequilibrium between parental alleles *j* and *l* in the pair of parental lines *P*_1_ and *P*_2_ which can be computed as 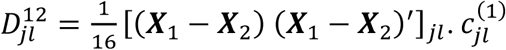 is the recombination frequency between parental alleles *j* and *l* in the parental lines obtained from the absolute genetic distance *d*_*jl*_ in Morgan as 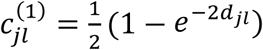 (Haldane 1919). When *j* and *l* refer to parental alleles at the same QTL, it holds 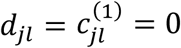. This formula given in [Eq. 3] can be applied analogously in every case presented in Figure 2: three-way crosses, backcrosses and two-way crosses. See File S1 for a detailed derivation of the covariance in DH-1 progeny [Eq. 3] and File S2 for an extension to DH progeny derived after selfing generations and to recombinant inbred lines at different selfing generations.

**Table 1.** Overview of genotypic covariance between loci *j* and *l* for different populations derived from the F1’ generation based on the disequilibrium parameter in pairs of parental lines.

| Population | Genotypic variance-covariance $\Sigma_{jl}$ |
| --- | --- |
| DH generation $k^a$ | $(1 - 2c_{jl}^{(k)}) \Phi_{2jl} + (1 - 2c_{jl}^{(k)} + c_{jl}^{(k-1)})(1 - 2c_{jl}^{(1)}) \Phi_{1jl}$ |
| RIL generation $k^b$ | $\left(1 - 2c_{jl}^{(k)} - \left(0.5(1 - 2c_{jl}^{(1)})\right)^k\right) \Phi_{2jl} + (1 - c_{jl}^{(k)})(1 - 2c_{jl}^{(1)}) \Phi_{1jl}$ |

<sup>a</sup> Doubled haploid (DH) lines derived after $k-1$ generations of selfing ( $k \in \mathbb{N}^*$ , $k = 1$ for DH lines derived directly from F1')
<sup>b</sup> Recombinant Inbred Lines (RIL) after $k$ generations of selfing ( $k \in \mathbb{N}^*$ , $k = 1$ for pseudo F2 generation)

### Indirect response to selection for parental contributions

We aim at predicting the full multivariate progeny distribution (mean, variance and pairwise covariances) for the agronomic trait, *P*_1_ genome-wide contribution (*C*) and *P*_1_ contribution at favorable alleles (*C*(+)). Therefore, we consider all three traits in the (4*p* × 3)-dimensional multi-trait effect matrix (***β***_*T*_ ***β***_*C*_ ***β***_*C*(+)_). Similarly as for one trait, the mean performance 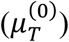 and mean genome-wide contribution of *P*_1_ in progeny before selection 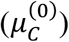 are derived as the mean of all four parents’ breeding values for each trait [Eq. 1]. As expected, 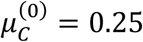 for four-way, three-way and backcrosses and 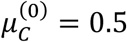 for two-way crosses. Progeny variances for all three traits are estimated using Eq. 2 and pairwise covariances in progeny are estimated as:

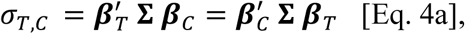

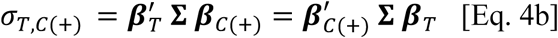

Progeny means and (co)-variances before selection can be used to estimate the expected response to selection on multiple traits. For this purpose, we used the Usefulness Criterion (Schnell and Utz 1975) in a multi-trait approach as illustrated in Figure 1. Assuming an intra-family selection of the progeny with the highest values for the agronomic trait with a selection intensity *i* and a selection accuracy of one (Figure 1A), the expected mean performance after selection 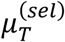 is defined as the usefulness criterion of the cross:

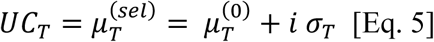

The correlated response to selection on *P*1 genome-wide contribution 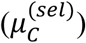 and *P*1 contribution at favorable alleles 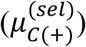 are (Falconer and Mackay 1996):

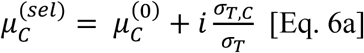

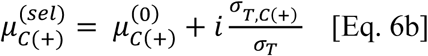

The contribution of *P*_1_ at unfavorable alleles after selection can be derived as:

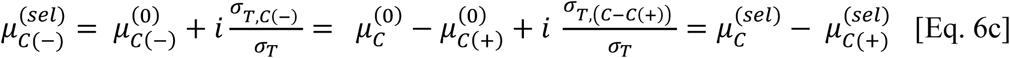

Figure 1C illustrates, in the case of a two-way cross (*P*_1_ x *P*_2_), the indirect response to selection on *P*_1_ genome-wide contribution 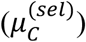 depending on the covariance *σ*_*T,C*_ that is mainly driven by the difference of performance between *P*_1_ and *P*_2_

### Simulation experiments

We performed two simulation experiments. The aim of the simulation experiment 1 was the validation of the presented formulas for the moments of the distribution of progeny from four-way crosses. In simulation experiment 2, we investigated different crossing schemes (two-way, three-way and backcrosses) in terms of genetic gain and donor contribution.

#### Genetic material

We considered 57 Iodent inbred lines from the Amaizing Dent panel (Rio *et al.* 2018). Iodent defines a heterotic group that has been derived 50 to 70 years ago and that is commonly used in maize breeding (Troyer 1999; Van Inghelandt *et al.* 2012). In the following we refer to these lines as elite lines. Elite lines were genotyped with the Illumina MaizeSNP50 BeadChip (Ganal *et al.* 2011). After quality control and imputation, 43,373 high-quality biallelic SNPs were retained. The genetic map was obtained by predicting genetic positions from physical positions (Jiao *et al.* 2017) using a spline-smoothing interpolating procedure described in Bauer *et al.* (2013) and the consensus dent genetic map in Giraud et al. (2014). We considered a quantitative agronomic trait (e.g. grain yield) implying *p* = 500 QTLs with known biallelic effects ***β***_0_ sampled from *N*(**0**_*p*_, 0.002 ***I***_*p*_).

### Simulation experiment 1: validation of UCPC

In order to validate the derivations for progeny (co)-variances and UCPC method in case of four-way crosses for DH and RIL progeny for selfing generations *k* ∈ [1, 6] (Table 1), we randomly generated 100 four-way crosses out of the 57 elite lines. For each cross, a set of 500 QTLs was randomly sampled among the 43,373 SNP markers across the genome to generate the agronomic trait. We also considered the first parent (i.e. *P*_1_) contributions: genome-wide (*C*) and at favorable alleles (*C*(+)). On one hand, we used algebraic formulas to predict the mean and (co)-variances for trait and contributions before selection within each cross (derivation). On the other hand, 50,000 DH or RIL progeny genotypes were simulated per cross at every selfing generation and the empirical mean and (co)-variances before selection were estimated (*in silico*). For *in silico* simulations, crossover positions were determined using recombination rates obtained with Haldane’s function (Haldane 1919). The correlated response to selection on *P*_1_ contributions after selecting the 5% upper fraction of progeny for the agronomic trait were either predicted using UCPC (derivation) or estimated after a threshold selection (*in silico*). The correspondence between predictors was assessed by the squared linear correlation and the mean squared difference between predicted (derivation) and empirical (*in silico*) values.

### Simulation experiment 2: evaluation of different multi-parental crossing schemes between donor and elite lines

We used UCPC to address the question of the best crossing scheme between a given genetic resource (donor *P*_1_, Figure 2), and elite lines. We identified the crossing scheme that maximized the short term expected genetic gain and evaluated donor genome contributions to the selected fraction of progeny. For this, we set up a simulation study where, at each iteration, an elite population of 25 lines was randomly sampled out of the 57 elite lines. Further, 500 QTLs were sampled among monomorphic and polymorphic markers in the elite population in order to conserve the frequency of monomorphic loci observed on 43,373 SNPs in the entire elite population. At each iteration, 100 intra-elite two-way crosses, backcrosses, and three-way crosses were randomly sampled as benchmark. Their progeny mean (μ_*T*_) and progeny standard deviation (*σ*_*T*_) for the agronomic trait were predicted by Eq. 1 and 2, respectively.

Within each iteration, 216 donor genotypes were constructed to cover a wide spectrum of donors in terms of performance and originality compared to the elite population. We defined three tuning parameters that reflect the proportions of six classes of QTLs (Dudley 1984) defined by the polymorphism between the donor and the elite population (Table 2). All possible combinations of the three tuning parameters varying from 0 to 1 with steps of 0.2 were considered. For each donor, we considered the simulated agronomic trait together with the donor genome contributions genome-wide (*C*) and at favorable alleles (*C*(+)). We defined the genetic gap with the elite population as the difference between donor and mean elite genetic values. The originality of the donor was defined as its mean pairwise modified Rogers distance (MRD) with elite lines.

**Table 2.**
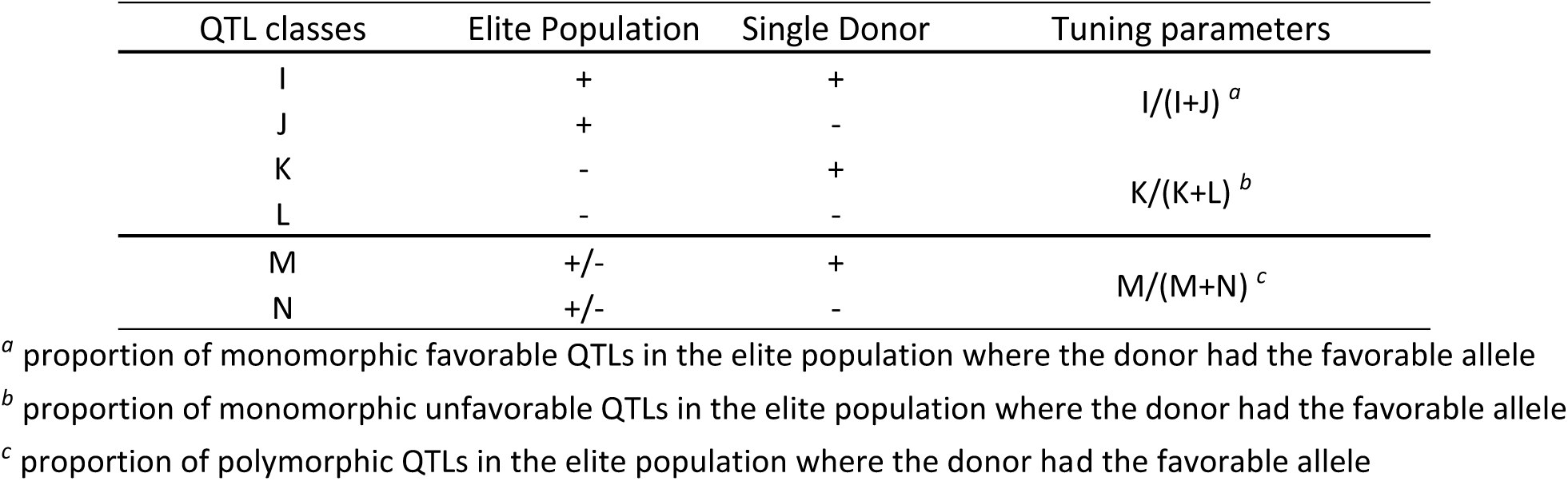
Classes of quantitative trait loci (QTL) and tuning parameters considered for simulating the donors. The favorable allele at QTL is denoted (+) and the unfavorable is denoted (-). A polymorphic QTL in the elite population is denoted (+/-).

For all possible 25 two-way crosses, 600 three-way crosses and 25 backcrosses between every donor and the elite population we predicted the progeny mean (μ) and the progeny standard deviation (*σ*) of each trait (Eq. 1 and Eq. 2) and the covariances between agronomic trait and contributions (*σ*_*T,C*_, Eq. 4a and *σ*_*T,C*(+)_, Eq. 4b). We defined the post-selection mean for the agronomic trait using Eq. 5 with selection intensity *i* corresponding to a selection pressure of 5%. For comparison between iterations, we subsequently standardized the UC for the agronomic trait based on the elite population by 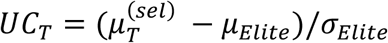, where μ_*Elite*_ is the mean and *σ*_*Elite*_ the genetic standard deviation of the elite population. After selection on the agronomic trait, the correlated response on donor contributions was estimated using Eq. 6 a-c. Finally, for each type of cross (two-way, three-way and backcrosses) and each donor, we identified the cross that maximized the expected genetic gain for the agronomic trait (*UC*_*T*_).

### Data availability

Simulations were based on genotypic maize data and genetic map deposited in File S4 at figshare. All simulations have been realized using R coding language (R Core Team 2017).

## RESULTS

### Simulation experiment 1: validation of UCPC

Predictions from the analytical derivations (Eq. 1, 2, 4a, 4b, 5, 6a, 6b) showed a high correspondence with empirical results from *in silico* simulations for the 100 DH-1 families (DH lines after *F1′*, Figure 2). The predicted progeny variance from derivations and from *in silico* simulations (Figure 3A-C) as well as the covariances between the agronomic trait and parent contributions (Figure 3D-E) showed squared correlations above 0.96. Predicted and simulated post-selection mean of the agronomic trait as well as predicted and simulated post-selection parental genome contributions showed correlations above 0.9 (Figure 3F-H) (R^2^ = 1.000 for Trait, R^2^ = 0.900 for *C* and R^2^ = 0.946 for *C*(+)). Validations for RIL and DH progeny derived from more selfing generations are presented in File S2.

**Figure 3.**
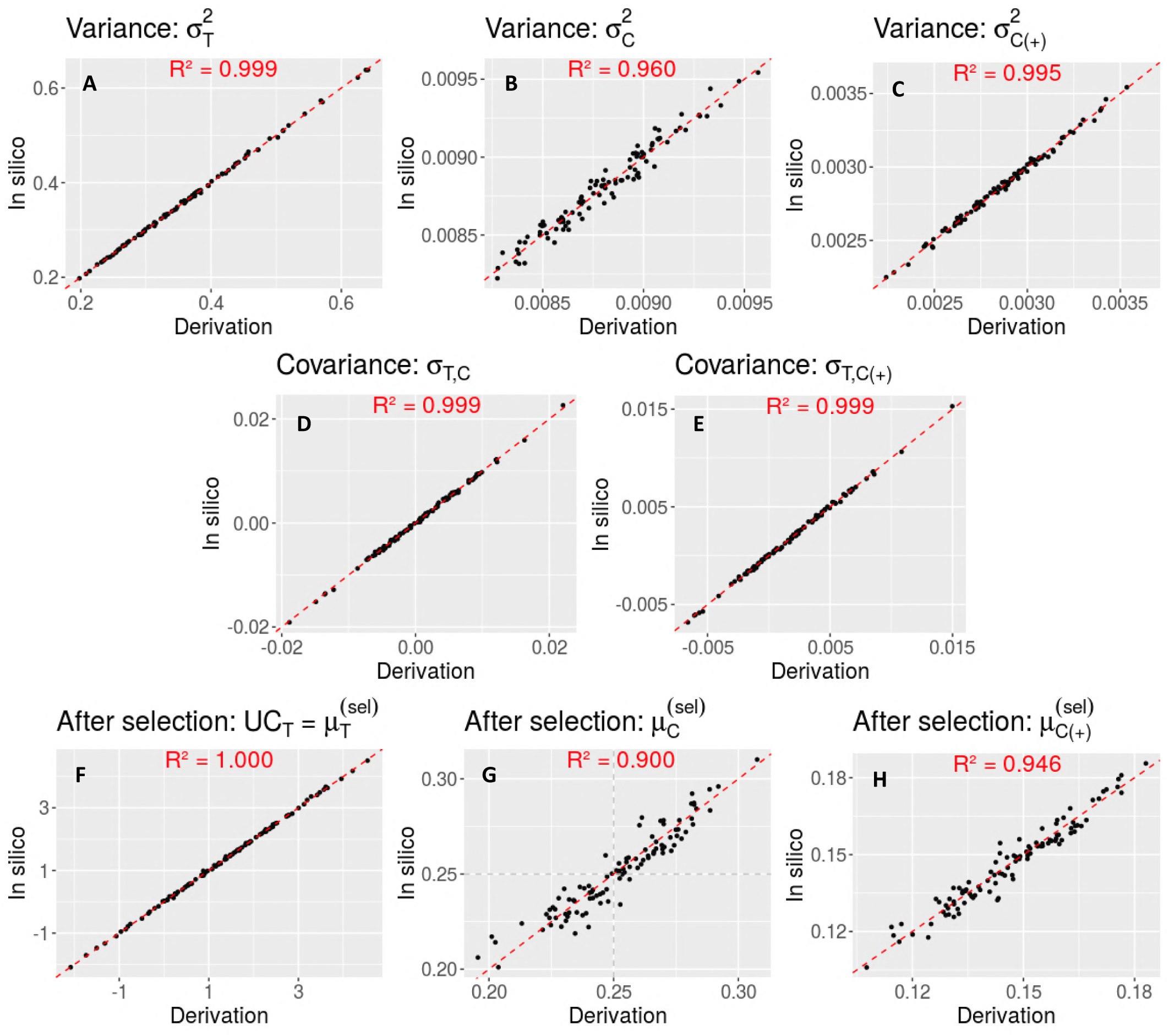
Comparison between predicted (derivation) and empirical (*in silico*) moments of the progeny distributions from 100 four-way crosses consisting of 50,000 DH-1 simulated progeny. Moments shown are (A) variance for the agronomic trail 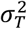, (B) variance for the genome-wide contribution 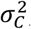, (C) variance of the contribution at favorable alleles 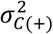, (D) covariance between agronomic trait and genome-wide contribution *σ*_*T,C*_, (E) covariance between agronomic trait and contribution at favorable alleles *σ*_*T,C*(+)_, (F) post-selection mean for the agronomic trait 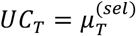, (G) post-selection mean for the genome-wide contribution 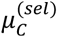, and (H) post-selection mean for the contribution at favorable alleles 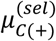. Squared correlations between predicted and empirical values are given within each plot.

### Simulation experiment 2

#### Intra-elite multi-parental crosses: a benchmark

Considering only the elite population generated at each iteration, the mean average performance over 20 iterations was μ_*Elite*_ = 0.067 ± 1.009 and the mean elite standard deviation was *σ*_*Elite*_ = 0.748 ± 0.107. We observed (Table 3) that intra-elite three-way crosses generated more progeny standard deviation (*σ*_*T*_) (0.576 ± 0.034) than two-way crosses (0.510 ± 0.026) and backcrosses (0.442 ± 0.022). In terms of progeny mean (μ_*T*_), differences were not significant between types of crosses. The gain in *σ*_*T*_ yielded a higher usefulness criterion (*UC*_*T* *mean*_) with three-way crosses (1.599 *σ*_*Elite*_ ± 0.317) than two-way crosses (1.461 *σ*_*Elite*_ ± 0.268). On the contrary, when only considering the best cross in terms of gain for the agronomic trait (*UC*_*T* *best*_), two-way crosses led to a higher UC (3.115 *σ*_*Elite*_ ± 0.362) than three-way crosses (2.876 *σ*_*Elite*_ ± 0.420) or backcrosses (2.804 *σ*_*Elite*_ ± 0.377).

**Table 3.**
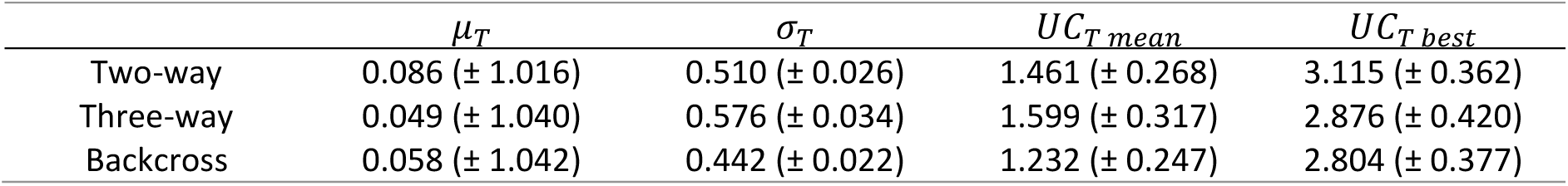
Intra-Elite crosses predicted progeny mean (μ_*T*_), progeny standard deviation (*σ*_*T*_) and resulting expected genetic gain *UC*_*T*_ with a selection pressure of 5%, once averaged over all crosses (*UC*_*T mean*_) and for the best cross identified (*UC*_*T best*_). For all parameters the mean (± standard deviation) over 20 iterations is given.

| | $\mu_T$ | $\sigma_T$ | $UC_{T\text{ mean}}$ | $UC_{T\text{ best}}$ |
| --- | --- | --- | --- | --- |
| Two-way | 0.086 ($\pm 1.016$) | 0.510 ($\pm 0.026$) | 1.461 ($\pm 0.268$) | 3.115 ($\pm 0.362$) |
| Three-way | 0.049 ($\pm 1.040$) | 0.576 ($\pm 0.034$) | 1.599 ($\pm 0.317$) | 2.876 ($\pm 0.420$) |
| Backcross | 0.058 ($\pm 1.042$) | 0.442 ($\pm 0.022$) | 1.232 ($\pm 0.247$) | 2.804 ($\pm 0.377$) |

#### Donor genome contribution in multi-parental crosses

For each simulated donor, we identified the two-way cross, three-way cross and backcross that maximized the UC for the agronomic trait (*UC*_*T*_). Those crosses are denoted as best crosses in the following. We analyzed the relationship between donor contributions to the selected progeny of the best crosses and the genetic gap between the donor and the mean elite population (Figure 4). The genome-wide contribution, the contribution at favorable alleles, and the contribution at unfavorable alleles are shown in Figures 4A, 4B and 4C, respectively. For a given donor, the genome-wide donor contribution after selection was higher in the best two-way crosses than in the best three-way crosses or backcrosses. For illustrative purposes, we differentiated five cases from the worst donor carrying only unfavorable alleles at QTLs (case 0) to the best donor carrying favorable alleles at all QTLs (case 4). Starting from case 0, the selection tended to eliminate most of the donor genome in progeny until a lower bound (Figure 4A, 27.1% for the best two-way cross, 6.7% for the best three-way cross and 6.3% for the best backcross). Very badly performing donors (case 1; genetic gap ≤ −5), i.e. carrying favorable alleles at maximum 180 QTLs, had little chance to pass their favorable alleles to the selected progeny (Figure 4B, 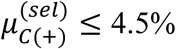 in the best two-way cross, 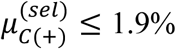 in the best three-way cross and 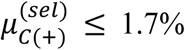 in the best backcross). When the performance of the donor increased (case 2; −5 < genetic gap ≤ 5), a higher portion of the donor genome was retained in the selected progeny (Figure 4B). With an increased number of favorable alleles (case 2), genome-wide donor contribution increased linearly with the genetic gap due to both, the selection of favorable alleles from the donor (Figure 4B) and the linkage drag with unfavorable alleles (Figure 4C). This linear trend continued until the donor had mainly favorable alleles (case 3; 5 < genetic gap). In case 3, we observed a linear increase of donor contribution at favorable alleles (Figure 4B). A correlated decrease of donor contribution at unfavorable alleles was observed at a nearly constant genome-wide contribution. Finally, in case 4, the genome-wide contribution was equal to an upper bound limit (Figure 4A, 72.6% for the best two-way cross, 42.9% for the best three-way cross and 43.5% for the best backcross).

**Figure 4.**
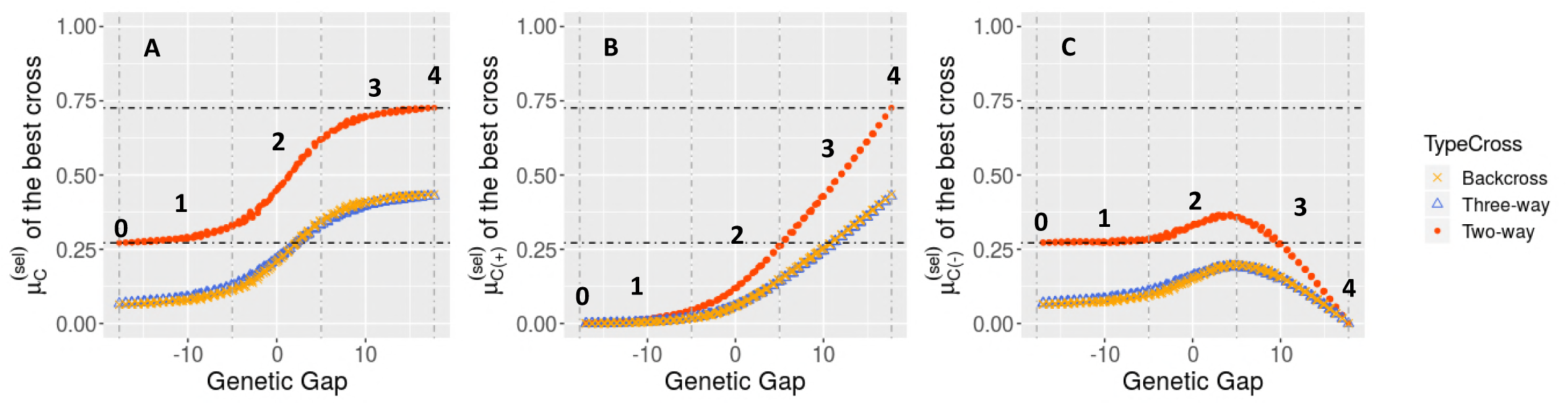
Donor contribution to the selected progeny of the best two-way cross (Donor*Elite), the best three-way cross ((Donor*Elite1)*Elite2) and the best backcross ((Donor*Elite1)*Elite1), depending on the genetic gap between donor line and the elite population. Each data point corresponds to the progeny of the best cross and is colored depending on the type of cross. (A) Donor genome-wide contribution after selection 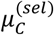, (B) donor genome contribution at favorable alleles after selection 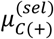 and (C) donor genome contribution at unfavorable alleles after selection 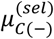. Numbers (0, 1, 2, 3, 4) correspond to illustrative cases based on genetic gap referred in the text. Illustrative cases 0 and 4 correspond to the worst and best donor respectively. Illustrative cases 1, 2, 3 are delimited by genetic gap values −5, 5 as represented by the vertical dashed lines.

#### Comparison of genetic gain among multi-parental crossing schemes

When the donor outperformed the elite population, the best two-way cross was more likely yielding a higher genetic gain than the best three-way cross or back-cross (Figure 5A). On the contrary, when the donor underperformed the elite population, the best three-way cross and backcross yielded a higher genetic gain than the best two-way cross. The higher progeny standard deviation (*σ*_*T*_) in the best two-way cross compared to the best three-way cross or backcross (Figure 5B) did not compensate the loss in progeny mean (μ_*T*_) (Figure 5C) in the best two-way cross. For a given genetic value of the donor, its originality did not impact the ranking between two-way crosses, three-way crosses and backcrosses (results not shown). A similar comparison between three-way crosses and backcrosses showed that the best backcross yielded similar μ_*T*_ (Figure 5B) but lower *σ*_*T*_ than the best three-way cross (Figure 5C), especially when the donor had a genetic value close to the best elite lines. This resulted in a slightly higher expected genetic gain in three-way crosses compared to backcrosses (Figure 5A).

**Figure 5.**
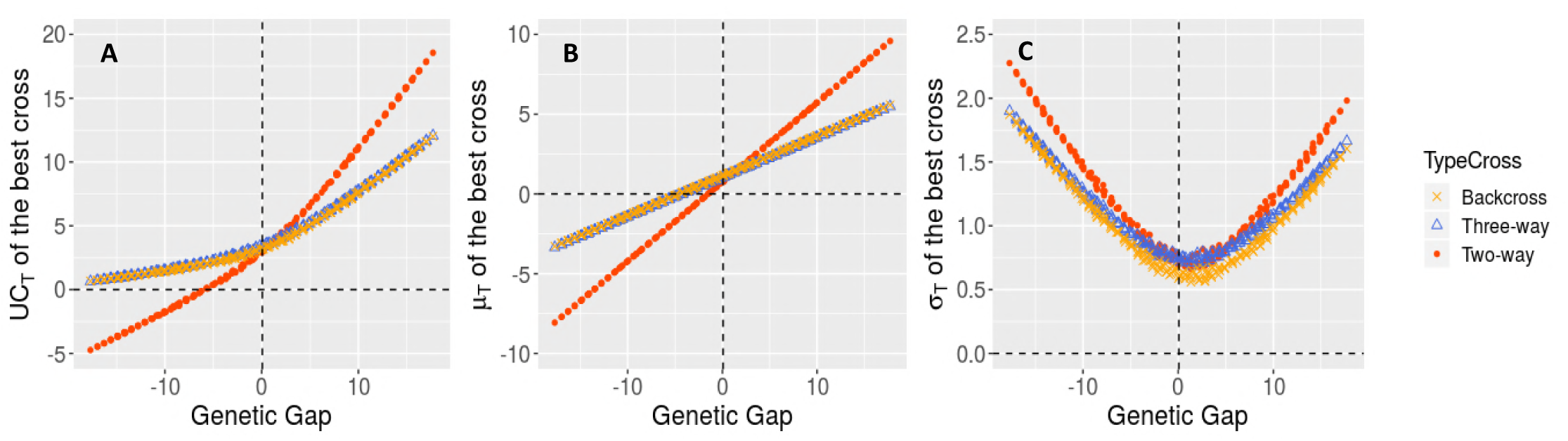
Comparison of the best two-way cross (Donor*Elite), the best three-way cross ((Donor*Elite1)*Elite2) and the best backcross ((Donor*Elite1)*Elite1), depending on the genetic gap (x-axis) with the elite population. Each data point corresponds to the progeny of the best cross and is colored depending of the type of cross. Comparison for the (A) expected genetic gain *UC*_*T*_, (B) progeny mean (μ_*T*_), and (C) progeny standard deviation (*σ*_*T*_).

## DISCUSSION

### Usefulness criterion for quantitative traits in multi-parental crosses

Accurate predictors of progeny variance accounting for the map position of loci and linkage phase of alleles in parents have been recently derived for biparental crosses (Lehermeier *et al.* 2017b; Osthushenrich *et al.* 2017). Nonetheless, breeders might use multi-parental crosses implying more than two parents to combine best alleles segregating in the breeding population. Therefore, we extended derivations given by Lehermeier *et al.* (2017b) for two-way crosses to four-way crosses by accounting for linkage disequilibrium between pairs of parental lines. We validated the derived genetic variance of RIL and DH progeny of four-way crosses by simulations (Figure 3, File S2). As expected, the formula for four-way crosses reduces to the one given by Lehermeier *et al.* (2017b) in case of two-way crosses (File S1). The results from our simulations showed that, considering elite material only, three-way crosses generate on average more variance than two-way crosses or backcrosses, resulting in higher genetic gain (Table 3). Nevertheless, the best possible cross (i.e. maximizing the expected genetic gain) was a two-way cross for most iterations (90%). This can be explained by the fact that crossing the two best elite lines generates more genetic gain than crossing them to a third less performant elite line, despite a potential gain in progeny variance. Notice that we considered only one polygenic agronomic trait but three-way crosses can be more advantageous for bringing complementary alleles for several traits. Under the formulated assumptions and with available marker effects (see discussion below), the general formula to predict mean and variance of four-way cross progeny makes it possible to identify the multi-parental cross that maximizes a given multi-trait selection objective (see discussion below) without requiring computationally intensive *in silico* simulations of progeny. The generalization to several generations of selfing for RIL progeny enables in addition to differentiate crosses releasing differently the variance in time (File S2). The presented formula for four-way crosses can also be applied to crosses of two heterozygous parents by considering its phased genotypes as four separate parents. Doing so, our approach can be adapted for heterozygous plant varieties that are common in perennial species and for crosses with hybrids, as well as for animal breeding where the prediction of Mendelian sampling variance can be very useful for mating decisions (Bonk *et al.* 2016).

### Parental contributions in multi-parental crosses under selection

Frisch and Melchinger (2007) derived the expected variance of parental contribution before selection in fully homozygote progeny accounting for linkage disequilibrium between loci assuming a biparental cross and considering only polymorphic loci. In this study, we proposed an original way to follow parental genome contribution to the selected fraction of progeny in multi-parental crosses, namely UCPC. It is grounded in a normal approximation of the probability mass function of parental contribution (Hill 1993; Frisch and Melchinger 2007) and progeny variance derivations. In the specific case of DH lines derived from two-way crosses or backcrosses and considering one chromosome of 100cM, our prediction of parental genome contribution variance converged to the one of Frisch and Melchinger (2007) when increasing the number of loci (File S3). However, the previous literature did not combine parental contributions with quantitative traits. Our original multivariate UCPC approach enables to predict the covariance between parental genome contributions and traits of economic interest. Based on multivariate selection theory, UCPC predicts the expected realized parental genome contribution after selection on traits of interest. It allows to follow parental genome contribution inheritance over generations and provides the likelihood of reaching a specific level of parental contribution while prescreening the most performing lines. Such information can guide breeders and researchers to determine the minimal number of progeny to derive from a cross between a donor and one or several elite lines so that the expected donor contribution after selection can reach a targeted value.

Predicted genome-wide donor contribution to progeny after selection was bounded to a minimum in case of the worst donor and a maximum in case of the best donor. In line with the predicted distribution of parental genome contribution before selection obtained in maize by Frisch and Melchinger (2007), these results show that in one selection cycle with a reasonable selection intensity (e.g. 5%) it is unlikely to get completely rid of unfavorable parental alleles. Parental genome contribution was bounded in selected progeny due to the low probability of combining all alleles from a single parent.

### Recommendations for donor by elites crosses

Using UCPC, we addressed the question of polygenic trait introgression from an inbred donor to inbred elite recipients with a focus on common plant breeding crossing schemes: two-way, three-way and backcrosses. We assumed that the objective was to derive in one selection cycle an inbred progeny that combined donor favorable alleles in a performing elite background. Such progeny can be used as parental lines for new crosses in order to quickly introgress new favorable alleles in a breeding program. Such a short term vision of genetic resource integration can be complementary to a longer term pre-breeding approach using exotic material (Bernardo 2009; Gorjanc *et al.* 2016; Yu *et al.* 2016). As expected, donors underperforming the elite population (inferior donor) yielded a higher genetic gain when complemented by two elite lines in three-way crosses or by twice an elite line in backcrosses rather than by a single elite line in two-way crosses. In this case, there is an advantage of crossing schemes involving, on average before selection, only one fourth of the donor genome instead of half of the donor genome as it would be the case for a two-way cross. On the contrary, two-way crosses were more adapted to donors outperforming the elite population. If the donor showed a similar performance level as the elite lines, no general rule could be drawn. In such a case, we recommend to identify the best crossing scheme by predicting every potential cross using the UCPC approach. As expected under a lower dilution of donor alleles into elite alleles in two-way crosses compared to three-way crosses or backcrosses, the predicted genome-wide donor contribution to selected progeny was higher in the best two-way cross than in the best three-way cross or the best backcross (Figure 4A).

We observed for a polygenic trait that, despite a lower competition between donor and elite favorable alleles, backcrosses were not significantly superior to three-way crosses for maintaining higher donor contribution at favorable alleles (Figure 4B). In addition, backcrosses generated less progeny variance (Figure 5C) but similar progeny mean than three-way crosses, resulting in a lower genetic gain (Figure 5A). This observation depends on the elite population considered. For instance, it might not hold if one unique elite line highly outperforms all other lines. More generally, while backcrosses only combine donor alleles with alleles of one elite parent, three-way crosses combine donor alleles with alleles of two complementary elite lines and are thus closer to material generated at the same time using two-way crosses in routine breeding. For these reasons, we suggest that three-way crosses should be preferred over backcrosses for polygenic trait introgression in elite germplasm. Our results support *a posteriori* the crossing strategy adopted in the Germplasm Enhancement of Maize project (GEM, e.g. Goodman 2000). In GEM, maize exotic material has been introgressed into maize elite private lines using three-way crosses implying two different private partners. With the possibility to efficiently predict the progeny distribution of three-way crosses (UCPC), the best crossing partners can be identified to meet the targeted outcome in short time which allows to fully profit of the advantages of three-way crosses.

### Multivariate selection for agronomic traits and parental contributions

We observed that badly performing donors had little chance to pass their favorable alleles to progeny selected for their agronomic trait performance. This is a consequence of the negative covariance between the performance for the trait and donor contribution in case of an inferior donor (Figure 1C). To prevent this loss of original alleles, we could account for such tension in the multivariate context, for instance by applying a truncation on donor contribution before selecting for the trait using the truncated multivariate normal theory (Horrace 2005) or vice versa. Otherwise, selection on donor contribution and the agronomic trait can be applied jointly by building a selection index, which is promising to balance short term genetic gain and long term genetic diversity (i.e. selection on donor contribution) according to specific pre-breeding strategies.

More generally, the multivariate context provides the opportunity to deal with several quantitative traits on which selection is directly or indirectly applied. Further traits for which genome-wide estimated marker effects or QTL effects are available can be considered. For external genetic resource utilization, it enables to introgress secondary traits such as polygenic tolerances to biotic or abiotic stresses (e.g. drought tolerance), while agronomic flaws (e.g. plant lodging) can be counter-selected using threshold selection. Recently it has been shown by Akdemir et al. (2018) how the improvement of multiple traits can be addressed with multi-objective optimized breeding strategies.

### Practical implementation of UCPC in breeding

In practice, marker effects estimated with whole-genome regression models can be used in lieu of QTL effects that are unknown. Such effects should be estimated on a proper training population mixing both elite lines and original genetic resources. Marker effects can be estimated using Bayesian Ridge Regression as suggested in Lehermeier *et al.* (2017a; b) to derive an unbiased estimator of progeny variance (PMV: posterior mean variance). In our simulation study, we considered only biallelic QTL effects. As we formulated a multi-allelic model, population-specific additive effects could be considered straightforwardly. Considering that the donor might have a different origin than the elite lines (e.g. other heterotic group in hybrid crops), it might be of interest to use parental specific effects estimated by e.g. multivariate QTL mapping (Giraud *et al.* 2014) or genome-wide prediction models (Lehermeier *et al.* 2015).

Our approach is totally generic and can deal with any information on the position and the effect of QTLs. However, main assumptions should be discussed at this point. We assumed known true genetic positions of QTLs and no interference during crossover formation to derive recombination frequencies (Haldane 1919). In practice, the precision of recombination frequency estimates is a function of the available mapping information and the frequency of interference. Furthermore, recombination frequency might vary among the same species (Bauer *et al.* 2013) impairing the accuracy of variance prediction. To limit this risk we suggest to use a multi-parental consensus map (e.g. Giraud *et al.* 2014). Furthermore, derivations assumed no selection before developing progeny. However, selecting progeny from which to derive DH lines is likely in practice. This can involve voluntary molecular prescreening for disease resistance (e.g. during selfing generations) or practical limitations (e.g. originating from low DH induction rates). If the genetic correlation between those traits and the traits considered within UCPC is null, the derived progeny distribution and UC for the four-way crosses will still hold.

The derived formula for progeny mean and variance holds for mono- and oligo-genic traits, whereas the usefulness criterion underlying UCPC uses normal distribution properties. When considering traits involving a sufficient number of underlying QTLs, as it is the case for most agronomic traits and parental genome contributions, this assumption of normality is likely guaranteed by the central limit theorem. If only a limited number of known QTLs should be introgressed from a donor, an allele pyramiding strategy will be more suitable (Hospital and Charcosset 1997; Charmet *et al.* 1999; Servin *et al.* 2004). Furthermore, the predicted cross value (PCV) as recently suggested by Han et al. (2017) can be applied in this context and could be extended to multi-parental crosses considering our derivation of progeny variance.

We presented an IBD definition of parental genome contributions using a multi-allelic approach. The multi-allelic coding yields covariance matrices that are four times larger compared to using a biallelic coding. In practice, to obtain a less computationally intensive solution, the genotyping matrix can be reduced to a bi-allelic coding which yields an identity by state (IBS) parental genome contribution that informs on the sequence similarity between one parent and progeny (see File S3). However, in such a case parental contributions do not sum up to one and it cannot be accounted for multi-allelic (i.e. haplotypic) effects. For biparental crosses (i.e. two-way and backcrosses), an IBS approach (File S3) considering only polymorphic markers homogeneously covering the genome can be used as an approximation of the IBD contribution.

### Future research directions

UCPC is opening several future research directions. We illustrated the use of UCPC for a simple donor introgression problem but it can be extended to more complex problematics commonplace in breeding. For instance, UCPC can be applied to evaluate the interest of introgressing several donors, e.g. evaluate the interest of combining alleles from two donors (*D*_1_ and *D*_2_) with elites (*E*_1_ and *E*_2_) in (*D*_1_ × *E*_1_) × (*D*_2_ × *E*_2_) or (*D*_1_ × *D*_2_) × (*E*_1_ × *E*_2_).

Mating design optimizations, i.e. finding an optimized list of crosses to realize each year, accounting for a compromise between short and long term genetic gain have been investigated using two-way crosses and parental means as predictor of the expected gain and the inbreeding rate in the next generation (Beukelaer *et al.* 2017; Gorjanc *et al.* 2018). Applying UCPC within the context of mating design optimization would enable to account for parental complementarity through the use of progeny variation, i.e. within cross variance, as proposed by Shepherd and Kinghorn (1998), Akdemir and Sánchez (2016) and Müller *et al.* (2018). Furthermore, UCPC would enable to use parental contribution to the selected fraction of progeny to predict the realized inbreeding in the next generation. We conjecture that considering the realized parental genome contribution together with the usefulness criterion in UCPC is promising for mating design optimization to manage short and long term genetic gain in breeding programs. Future research will also be needed to investigate the use of multi-parental crosses in mating design optimizations. Hereby, UCPC that efficiently predicts the progeny distribution of crosses with up to four parents will represent a good starting point for further research.

### Conclusions

We developed, validated and illustrated the usefulness criterion parental contribution (UCPC) that evaluates the interest of multi-parental crosses based on the expected genetic gain (UC) and the parental contributions (PC) in the next generation. UCPC allows to (i) predict the progeny variance of four-way crosses accounting for linkage disequilibrium and to (ii) follow all parental genome contributions to the selected progeny to evaluate the interest of a cross regarding an objective that is a function of the expected performance and the diversity in the selected progeny. Illustration of the use of UCPC in the context of polygenic trait introgression from a donor to elite recipients enabled to draw some major recommendations. As expected, three-way crosses and backcrosses were more adapted to donors underperforming the elite population (inferior donor) while two-way crosses were more adapted to donors outperforming the elite population. We also suggested that three-way crosses should be preferred over backcrosses for polygenic traits introgression. Furthermore, we highlighted the importance of a compromise between UC and PC in case of an inferior donor.

## ACKNOWLEDGEMENTS

The authors thank the Amaizing program for genotypes used in simulations. This research was funded by RAGT 2n and the ANRT CIFRE Grant n° 2016/1281 for AA.

## File S1

### Derivation of linkage disequilibrium parameter in progeny for four-way cross and specific case of two-way cross, three-way cross and backcross

Here we derive the linkage disequilibrium parameter of doubled haploid progeny derived from the *F*_1_’ generation of a four-way cross (Figure 1 S1), while we give an extension for DH lines generated from higher selfing generations and for recombinant inbred lines in File S2. The crossing scheme for a four-way cross visualizing parental and potential progeny haplotypes is given in Figure 1 S1. Gametes from a four-way cross with four different parents (P1, P2, P3, and P4) correspond to gametes from six biparental crosses (P1xP2, P3xP4, P1xP3, P1xP4, P2xP3, P2xP4).

**Figure 1 S1.**
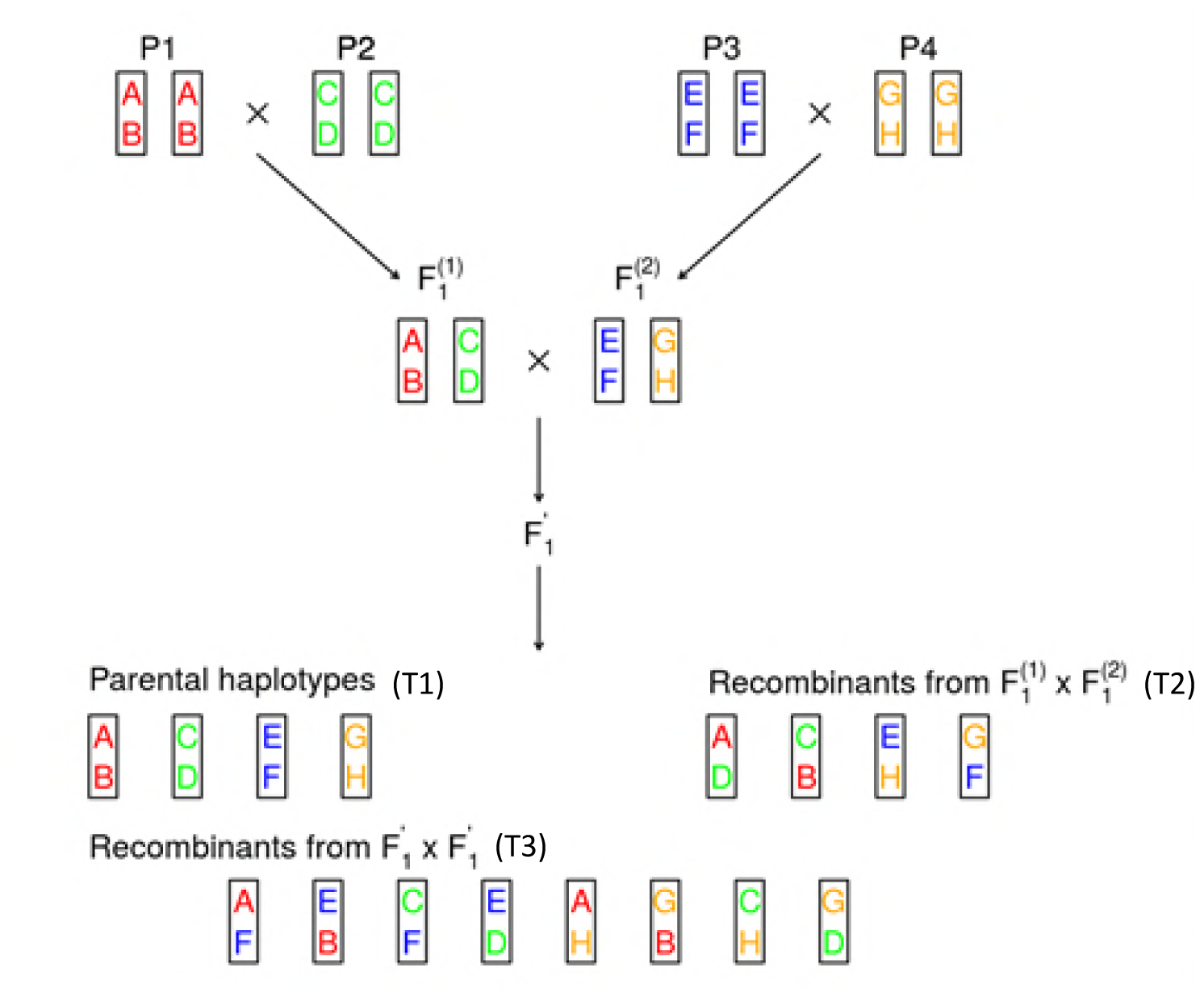
Visualization of crossing scheme and two-locus parental as well as progeny haplotypes of a four-way cross from parents P1, P2, P3, and P4. Potential types of haplotypes are denoted with T1, T2, and T3.

To derive the entries of the Linkage Disequilibrium (LD) matrix ***D*** of the progeny of the four-way cross, we derive the frequencies of all different possible haplotypes. For this, three types of haplotypes can be differentiated (namely, T1, T2 and T3).

The first type T1 corresponds to parental haplotypes, for example AB from Figure 1 S1. The frequency of the haplotype AB in the parents is:

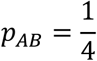

The frequency of AB in gametes from the cross 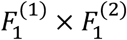 is:

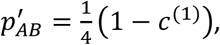

with *c*^(1)^ the recombination frequency and (1 - *c*^(1)^) the frequency that no recombination takes place within the cross 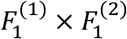.

Similarly, the frequency of AB in gametes from the cross 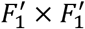 is:

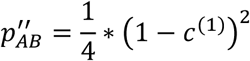

As there are four different parental haplotypes, the frequency of the type T1 haplotypes is:

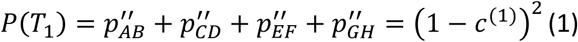

The second type T2 corresponds to haplotypes formed by recombination in the cross 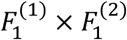, for example AD. The frequency of this haplotype in the parents is

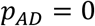

The frequency of AD in gametes from the cross 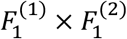 is:

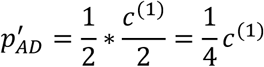

As 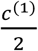 is the frequency of recombinants within 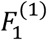, the frequency in the whole cross is reduced by a factor of 1/2. The frequency of AD in gametes from the cross 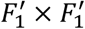 is:

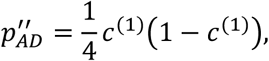

with (1 - *c*^(1)^) the frequency that no recombination takes place within the cross 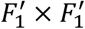.

Overall, the frequency of the type T2 haplotypes is:

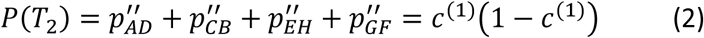

The third type T3 corresponds to haplotypes formed by recombination in the cross 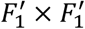, for example AF. The frequency of these haplotypes in the parents is:

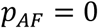

The frequency of AF in gametes from the cross 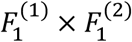 is:

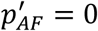

The frequency of AF in gametes from the cross 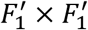 can be calculated as:

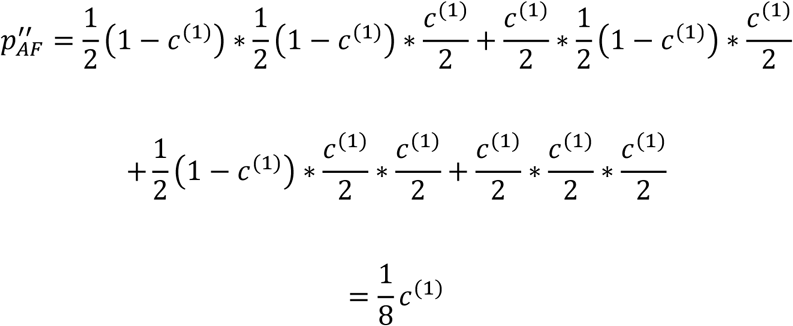

Overall, the frequency of the type T3 haplotypes is:

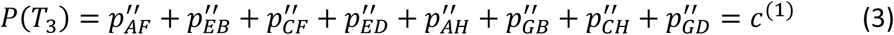

All the different haplotypes and frequencies are summarized in Table 1 S1.

We define *h*_*jl*_ = (*h*_*j*_, *h*_*l*_) a haplotype including loci *j* and *l*, with *h*_*j*_ and *h*_*l*_ the alleles of the haplotype at loci *j* and *l, h*_*j*_, *h*_*l*_ ∈ {0,1}. Using the frequencies of the three types of haplotypes, we derive the LD in the progeny between locus *j* and *l* as:

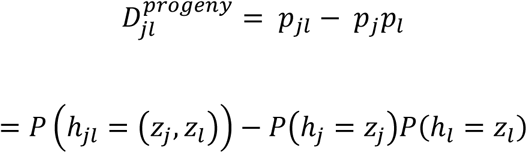

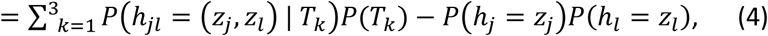

where *z*_*j*_ and *z*_*l*_ denotes realizations of *h*_*j*_ and *h*_*l*_.respectively.

For the conditional haplotype probabilities it holds:

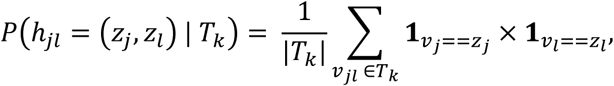

with |*T*_*k*_| the number of haplotypes of type *k, v*_*jl*_ = (*v*_*j*_, *v*_*l*_) a haplotype of type *k*, **1**_*vj==zl*_ (**1**_*vj==zl*_) an indicator equal to 1 if *v*_*j*_ = *z*_*j*_ (*v*_*l*_ = *z*_*l*_) and 0 otherwise.

For the allele frequencies it holds:

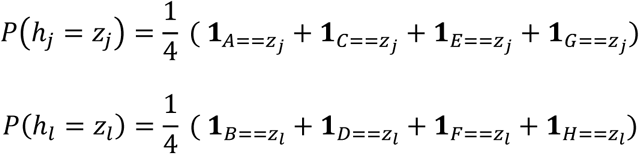

**Table 1 S1.** Different haplotype types, their frequency in the parents (G0), after the first cross (G1), after the second cross (G2) and the Linkage Disequilibrium (LD) in G2.

| Type | G0 | G1 | G2 | LD |
| --- | --- | --- | --- | --- |
| T1<sup>a</sup> | $\frac{1}{4}$ | $\frac{1}{4}(1 - c^{(1)})$ | $\frac{1}{4} * (1 - c^{(1)})^2$ | $\frac{1}{4} * (1 - c^{(1)})^2 - \frac{1}{16}$ |
| T2<sup>b</sup> | 0 | $\frac{1}{4}c^{(1)}$ | $\frac{1}{4}c^{(1)} * (1 - c^{(1)})$ | $\frac{1}{4}c^{(1)} * (1 - c^{(1)}) - \frac{1}{16}$ |
| T3<sup>c</sup> | 0 | 0 | $\frac{1}{8}c^{(1)}$ | $\frac{1}{8}c^{(1)} - \frac{1}{16}$ |

<sup>a</sup> Haplotypes: AB, CD, EF, GH (parental haplotypes)
<sup>b</sup> Haplotypes: AD, BC, EH, FG (recombinant from $F_1^{(1)} \times F_1^{(2)}$ )
<sup>c</sup> Haplotypes: AF, AH, CF, CH, EB, ED, GB, GD (recombinant from $F_1' \times F_1'$ )

Further, we use the linkage disequilibrium among two parents between loci *j* and *l*, which is exemplified for parent 1 and 2:

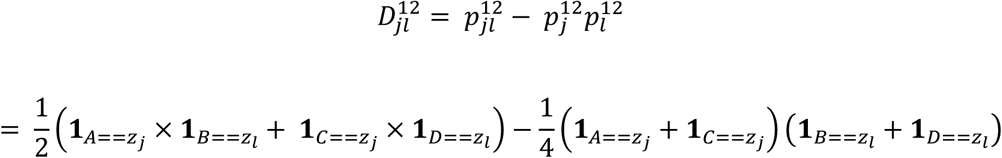

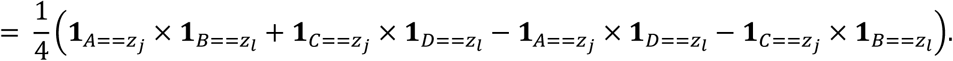

For sake of clarity, we abbreviate in the following **1**_*A==zj*_ with **1**_*A*_, **1**_*A==zl*_ with **1**_*B*_ and accordingly for the rest (C, D, E, F, G, H). Then we can reform the LD in the progeny as a function of the recombination 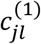 and the LD among two parents between loci *j* and *l*:

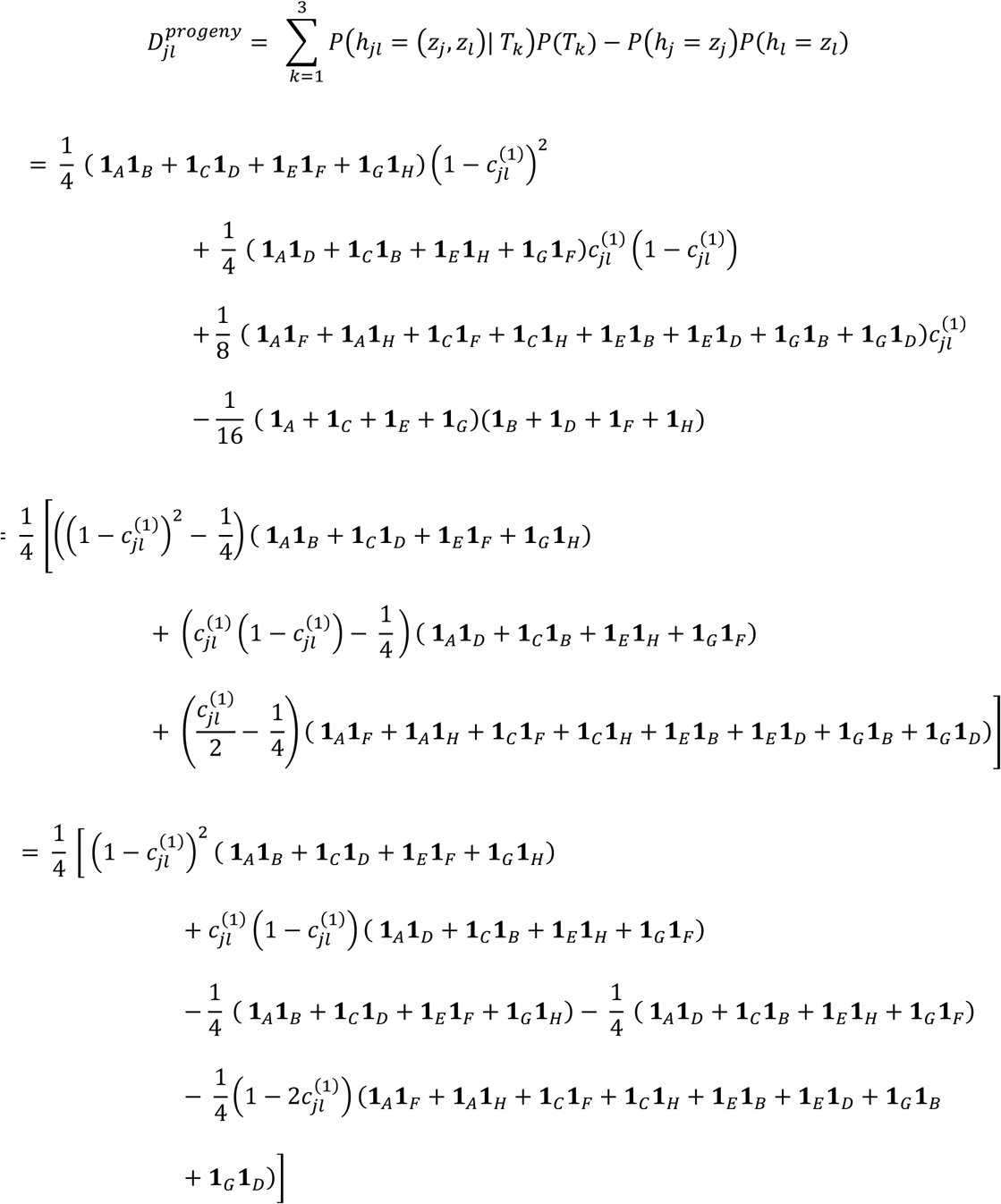

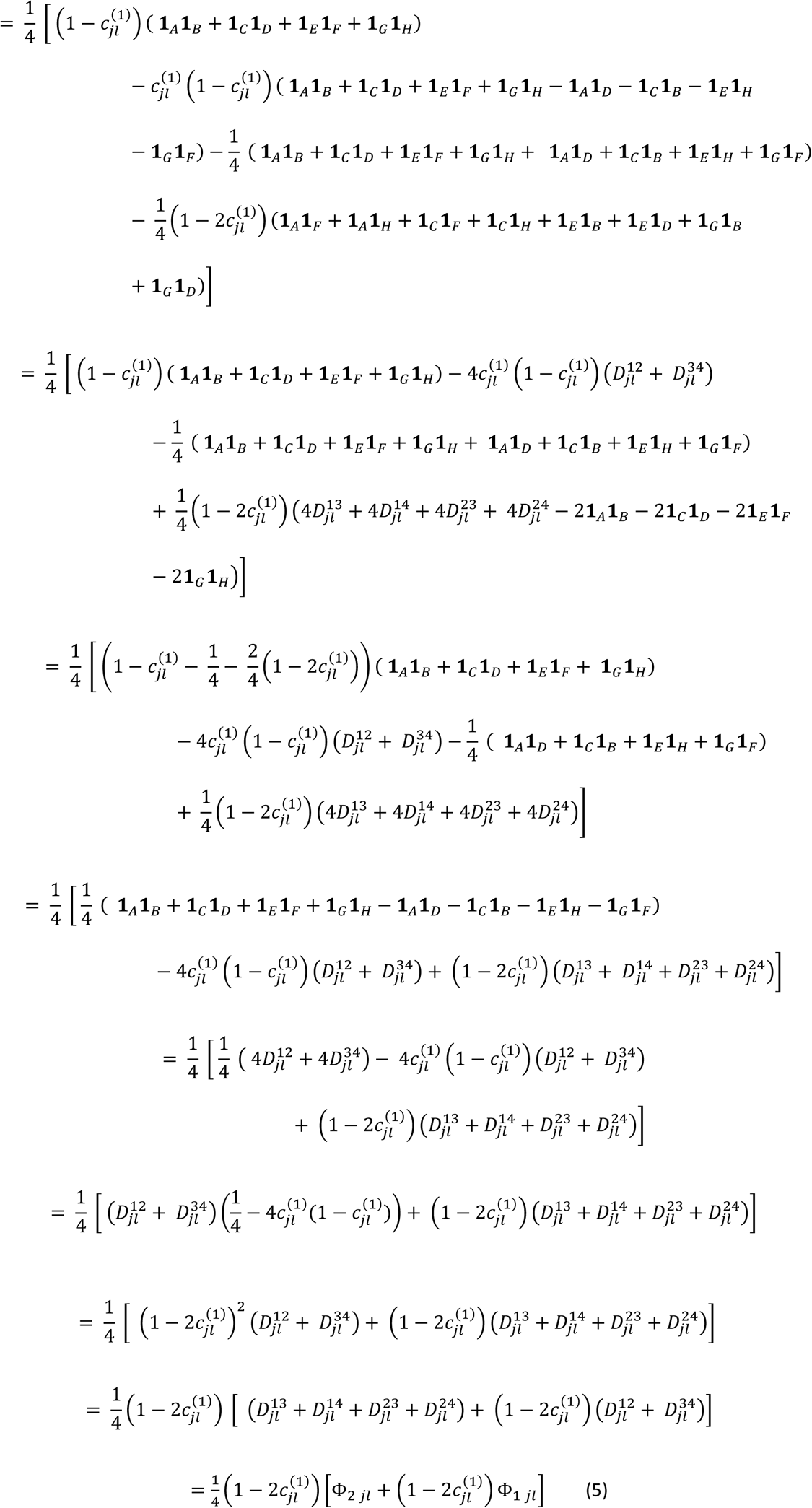

with 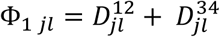 summing the LD values among parents that can be considered to be involved as biparental crosses in 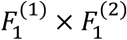 and with 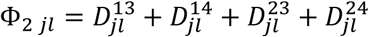 summing the LD values among parents that can be considered to be involved as biparental crosses in 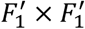.

The linkage disequilibrium parameter Φ_1_and Φ_2_ and equation (5) can be simplified in the case of two-way, three-way and backcrosses (Table 2 S1). For two-way crosses we arrive at the same variance covariance matrix elements Σ_*jl*_ as given by Lehermeier et al. (2017).

**Table 2 S1.**
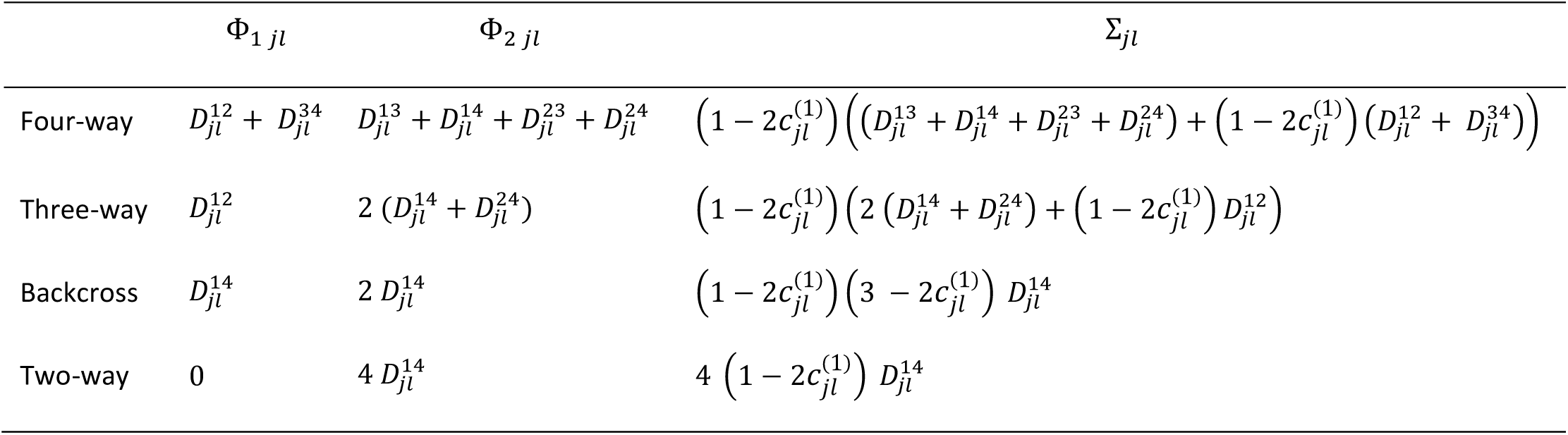
Linkage disequilibrium parameter between QTLs *j* and *l* in pairs of parental lines depending on the mating design.

| | $\Phi_{1\ jl}$ | $\Phi_{2\ jl}$ | $\Sigma_{jl}$ |
| --- | --- | --- | --- |
| Four-way | $D_{jl}^{12} + D_{jl}^{34}$ | $D_{jl}^{13} + D_{jl}^{14} + D_{jl}^{23} + D_{jl}^{24}$ | $\left(1 - 2c_{jl}^{(1)}\right) \left( (D_{jl}^{13} + D_{jl}^{14} + D_{jl}^{23} + D_{jl}^{24}) + \left(1 - 2c_{jl}^{(1)}\right) (D_{jl}^{12} + D_{jl}^{34}) \right)$ |
| Three-way | $D_{jl}^{12}$ | $2 (D_{jl}^{14} + D_{jl}^{24})$ | $\left(1 - 2c_{jl}^{(1)}\right) \left( 2 (D_{jl}^{14} + D_{jl}^{24}) + \left(1 - 2c_{jl}^{(1)}\right) D_{jl}^{12} \right)$ |
| Backcross | $D_{jl}^{14}$ | $2 D_{jl}^{14}$ | $\left(1 - 2c_{jl}^{(1)}\right) \left( 3 - 2c_{jl}^{(1)} \right) D_{jl}^{14}$ |
| Two-way | 0 | $4 D_{jl}^{14}$ | $4 \left(1 - 2c_{jl}^{(1)}\right) D_{jl}^{14}$ |

## File S2

### Validation of four-way cross formulas for DH-*k* and RIL-*k* and evolution of RIL variance depending on selfing generations

In File S1, we considered DH lines generated from F1’ (DH-1), i.e., only two meioses took place. Progeny variance for DH-1 is expressed in terms of parental expected recombination frequency *c*^(1)^ (Table 2 S1). For recombinant inbred lines (RILs) or when DH lines are generated from higher selfing generations, the expected frequency of recombinants increases depending on the number of selfing generations. In the following *k* denotes the generation from which progeny are derived (Figure 1). The expected frequency of recombinants in generation *k* can be derived from the genotype probabilities given in Broman (2012) as done in File S1 of Lehermeier et al. (2017). Hence, for DH lines after *k* generations, *c*^(1)^ in Table 2 S1 should then be replaced by *c*^(*k*)^, leading to the general four-way DH-*k* formula as shown in Table 1:

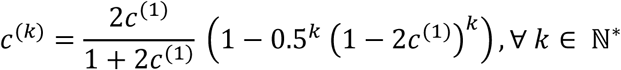

In case of RILs, no doubling of gametes takes place and the covariance for RILs after generation *k* is obtained by updating *c*^(*k*)^ by *c*^(*k*)^ + 0.5 [0.5(1 - 2*c*^(1)^]^*k*^, ∀ *k* ∈ ℕ^*^ (Table 1). Note that the variance-covariance of DH-*k* and RIL-*k* converge with increasing *k*.

Formulas for DH-*k* and RIL-*k* in the general case of four-way crosses have been validated by simulations for *k* ∈ ⟦1,6⟧ (Table 1 S2 and Table 2 S2). The observed high positive correlations (Table 1 S2) and low mean squared differences (Table 2 S2) between predicted (derivation) and empirical (*in silico*) values validate the presented formulas. Lower squared correlations between predicted and empirical values were observed for 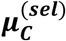 and 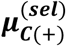 compared to the variances and covariances. This can be explained by sampling bias in *in silico* simulations (50,000 progenies) where the *P*_1_ parental genome contribution before selection slightly differed from the expected value of 0.25 for four way crosses (ranging from 0.249 to 0.251).

**Table 1 S2.** Squared correlations (R^2^) between empirical values (*in silico*) and predictions (derivation) per generation and type of progeny.

| Generation | $\sigma_T^2$ | $\sigma_C^2$ | $\sigma_{C(+)}^2$ | $\sigma_{T-C}$ | $\sigma_{T-C(+)}$ | $UC_T$ | $\mu_C^{(sel)}$ | $\mu_{C(+)}^{(sel)}$ |
| --- | --- | --- | --- | --- | --- | --- | --- | --- |
| DH1 | 0.999 | 0.960 | 0.995 | 0.999 | 0.999 | 1.000 | 0.900 | 0.946 |
| DH2 | 0.999 | 0.964 | 0.995 | 0.998 | 0.998 | 1.000 | 0.909 | 0.952 |
| DH3 | 0.999 | 0.966 | 0.995 | 0.999 | 0.999 | 1.000 | 0.914 | 0.955 |
| DH4 | 0.999 | 0.969 | 0.995 | 0.999 | 0.999 | 1.000 | 0.912 | 0.955 |
| DH5 | 0.999 | 0.961 | 0.994 | 0.998 | 0.998 | 1.000 | 0.914 | 0.955 |
| DH6 | 0.999 | 0.963 | 0.994 | 0.998 | 0.998 | 1.000 | 0.913 | 0.955 |
| RIL1 | 0.999 | 0.957 | 0.994 | 0.999 | 0.999 | 1.000 | 0.938 | 0.967 |
| RIL2 | 0.999 | 0.957 | 0.994 | 0.999 | 0.999 | 1.000 | 0.917 | 0.957 |
| RIL3 | 0.999 | 0.960 | 0.994 | 0.998 | 0.998 | 1.000 | 0.918 | 0.958 |
| RIL4 | 0.999 | 0.962 | 0.994 | 0.998 | 0.998 | 1.000 | 0.915 | 0.956 |
| RIL5 | 0.999 | 0.962 | 0.994 | 0.998 | 0.998 | 1.000 | 0.912 | 0.955 |
| RIL6 | 0.999 | 0.962 | 0.994 | 0.999 | 0.998 | 1.000 | 0.911 | 0.954 |

**Table 2 S2.** Mean squared difference between empirical values (*in silico*) and predictions (derivation) per generation and type of progeny.

| Generation | $\sigma_T^2$ | $\sigma_C^2$ | $\sigma_{C(+)}^2$ | $\sigma_{T-C}$ | $\sigma_{T-C(+)}$ | $UC_T$ | $\mu_C^{(sel)}$ | $\mu_{C(+)}^{(sel)}$ |
| --- | --- | --- | --- | --- | --- | --- | --- | --- |
| DH1 | 5.20E-06 | 3.28E-09 | 3.52E-10 | 5.99E-08 | 2.07E-08 | 8.44E-04 | 4.92E-05 | 1.42E-05 |
| DH2 | 5.09E-06 | 2.81E-09 | 3.16E-10 | 6.65E-08 | 2.24E-08 | 7.02E-04 | 3.83E-05 | 1.12E-05 |
| DH3 | 5.36E-06 | 2.56E-09 | 2.97E-10 | 4.74E-08 | 1.51E-08 | 6.49E-04 | 3.50E-05 | 1.03E-05 |
| DH4 | 4.56E-06 | 2.30E-09 | 2.87E-10 | 5.16E-08 | 1.66E-08 | 6.85E-04 | 3.55E-05 | 1.05E-05 |
| DH5 | 4.83E-06 | 2.88E-09 | 3.32E-10 | 5.95E-08 | 1.99E-08 | 6.40E-04 | 3.47E-05 | 1.03E-05 |
| DH6 | 4.76E-06 | 2.74E-09 | 3.14E-10 | 6.08E-08 | 1.96E-08 | 6.77E-04 | 3.47E-05 | 1.04E-05 |
| RIL1 | 2.25E-06 | 1.56E-09 | 1.81E-10 | 2.96E-08 | 9.80E-09 | 4.30E-04 | 2.51E-05 | 7.54E-06 |
| RIL2 | 3.26E-06 | 2.29E-09 | 2.69E-10 | 4.09E-08 | 1.37E-08 | 5.73E-04 | 3.40E-05 | 1.00E-05 |
| RIL3 | 3.93E-06 | 2.58E-09 | 3.05E-10 | 5.28E-08 | 1.72E-08 | 6.22E-04 | 3.34E-05 | 9.84E-06 |
| RIL4 | 4.49E-06 | 2.59E-09 | 3.02E-10 | 5.64E-08 | 1.81E-08 | 6.59E-04 | 3.43E-05 | 1.01E-05 |
| RIL5 | 4.91E-06 | 2.69E-09 | 3.10E-10 | 5.59E-08 | 1.83E-08 | 6.65E-04 | 3.53E-05 | 1.04E-05 |
| RIL6 | 4.91E-06 | 2.71E-09 | 3.13E-10 | 5.54E-08 | 1.83E-08 | 6.63E-04 | 3.59E-05 | 1.06E-05 |

Predicted RIL progeny variance for the simulated agronomic trait increased with the number of selfing generations considered (*k*) and converged toward DH progeny variance after five generations of selfing (*k* = 5) (Figure 1 S2). We observed that some crosses profited more from an increase in selfing generations by generating more variance compared to others. An example with two crosses is shown in Figure 2 S2. While the cross visualized in blue showed a higher variance in generation RIL-1 than the cross visualized in orange, it reached a plateau faster and showed a lower variance than the orange cross with *k* ≥ 3. Differences in the speed to release variance between crosses is likely due to differences in the recombination frequency between segregating QTLs in parental lines. This underlines the interest of predicting RIL progeny variance using proposed algebraic formula.

**Figure 1 S2.**
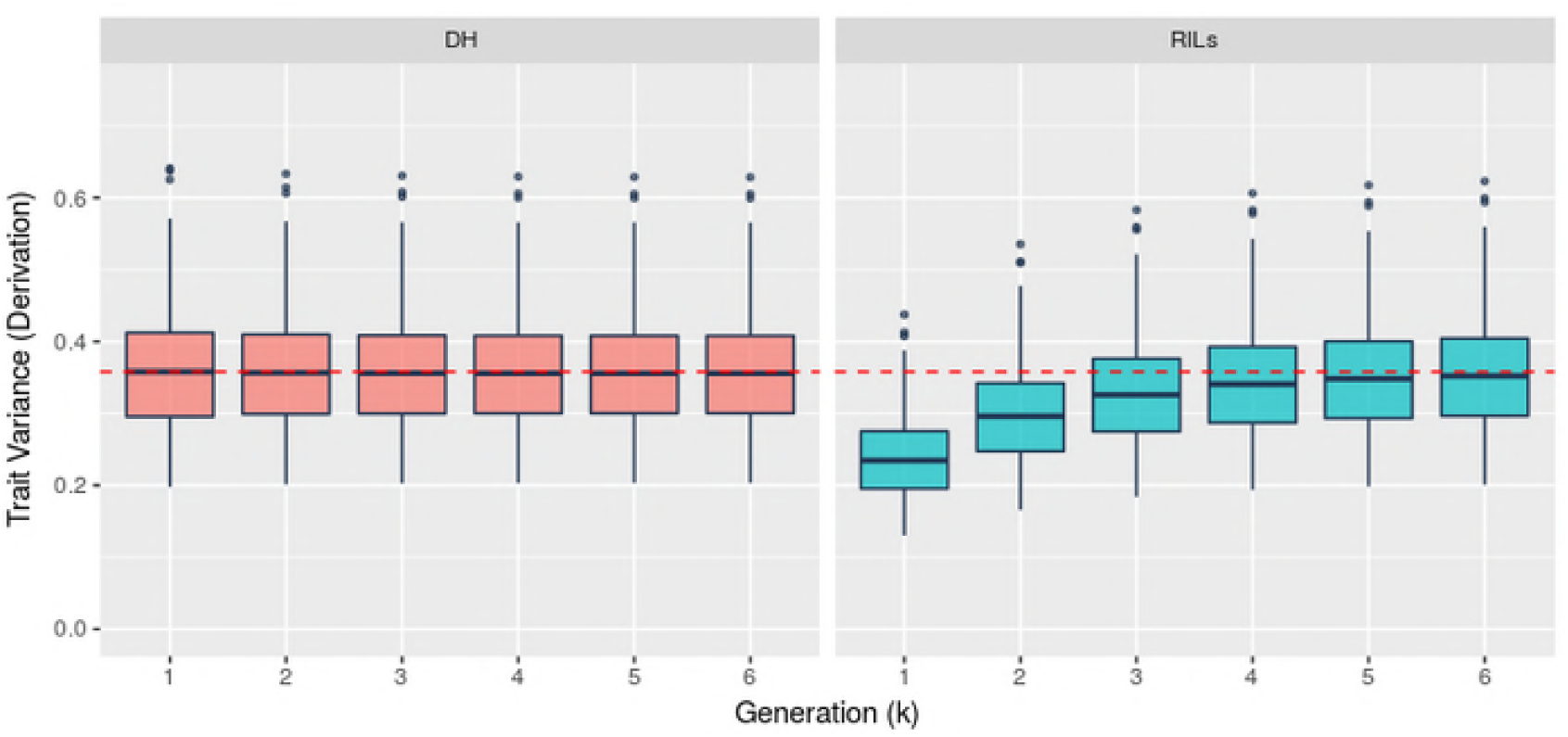
Evolution of predicted progeny trait variance depending on progeny type (DH, left or RIL, right) and generation (*k*). The red dotted line presents the median DH progeny variance over 100 crosses.

**Figure 2 S2.**
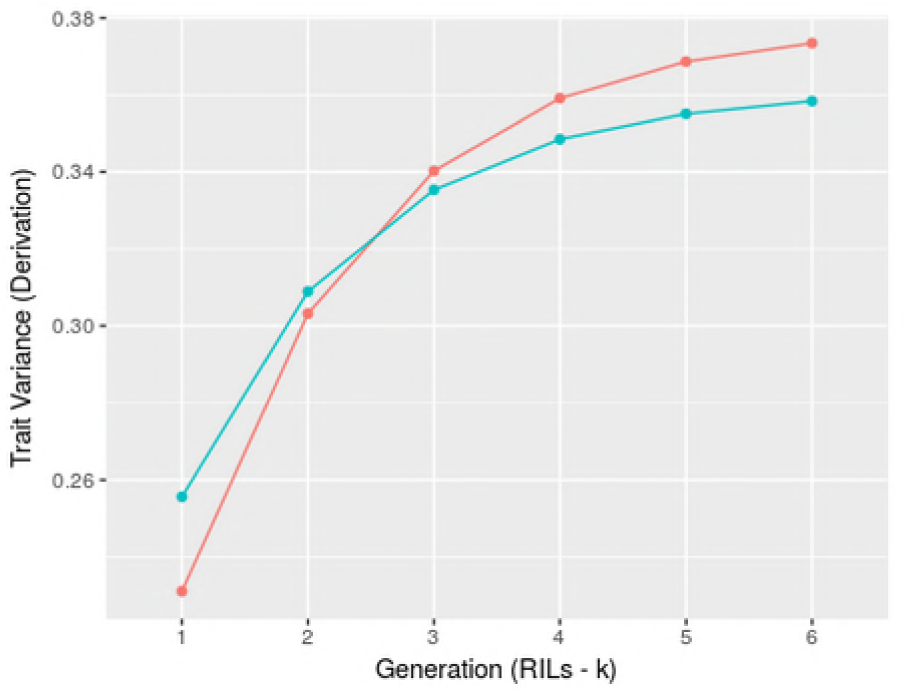
Example of two crosses showing different evolutions of predicted RIL progeny variance depending on the selfing generation (*k*).

## File S3

### Comparison of IBD parental contribution variance with Frisch and Melchinger (2007) and simplification to IBS contribution

We used an algebraic formula to predict the variance of *P*_1_ genome contribution in doubled haploid progeny derived from F1’ plants. We considered two-way crosses DH-1 (called (F1)-DH) and backcrosses DH-1 (called (BC1)-DH) and compared our results with the results given by Frisch and Melchinger (2007). We considered one chromosome of 100cM for which Frisch and Melchinger (2007) derived a variance of parental contribution of 0.1419 for (F1)-DH and 0.0945 for (BC1)-DH. We varied the number of loci *p* used in our approach and for each, we ran ten independent samplings of loci. We observed that the results from our approach converged with increasing number of loci to the solution given by Frisch and Melchinger (2007) (Figure 1 S3).

**Figure 1 S3.**
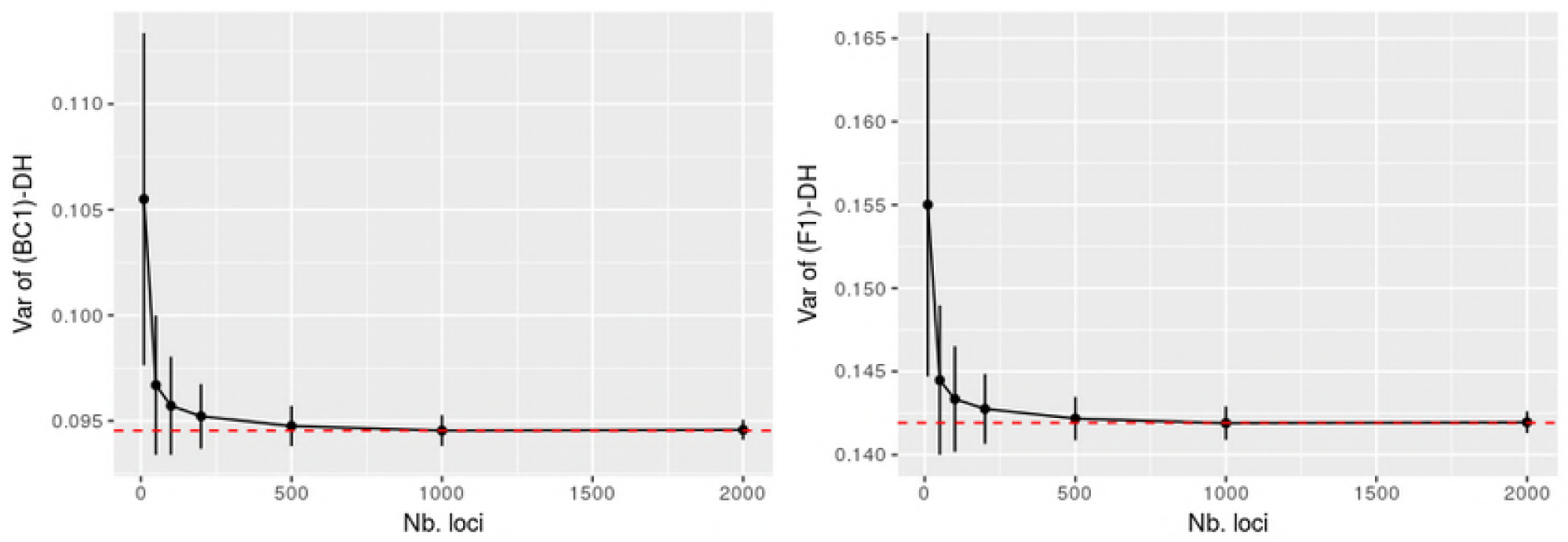
Average parental genome contribution variance (black dots) for (BC1)-DH (left) and (F1)-DH (right) from ten simulation replications (+/- standard deviation represented by black vertical lines) with different number of considered loci. Red dotted line shows the results given by Frisch and Melchinger (2007).

In cases where the origin of the allele is not of interest and an identical by state (IBS) similarity between progeny and parental lines is sufficient, the multi-allelic coding can be simplified to a biallelic coding. This reduces the size of the covariance matrix from (4*p* × 4*p*) to (*p* × *p*), with *p* being the number of loci considered. For this, let us define the genotyping matrix of parental lines in biallelic coding:

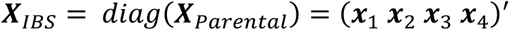

where, ***X***_*IBS*_ is a (4 × *p*)-dimensional matrix of genotypes. The (*p* × 4)-dimensional matrix of global parental contribution marker effects for each of the four parents can be defined as:

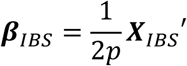

where, ∀ *i* ∈ [1; 4] ***β***_*IBS*_(., *i*) is the *p*-dimensional vector of marker effect to follow the IBS contribution of parent *i* and *p* is the total number of loci considered.

We denote the (*N* × *p*)-dimensional genotyping matrix of *N* doubled haploid (DH) progeny as ***X***_*IBS*-*Progeny*_ with element ***X***_*IBS*-*Progeny*_ (*j, l*), ∀ *j* ∈ [1, *N*], *l* ∈ [1, *p*] the genotype of progeny *j* at locus *l* coded as −1, 1 for the genotypes aa, AA, respectively. It results in the following (*N* × 4)-dimensional matrix of parental IBS contribution to progeny:

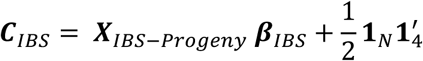

where, ∀ *j* ∈ [1; *N*], ∀ *i* ∈ [1; 4], ***C***_*IBS*_(*j, i*) is the parental line *i* contribution to progeny line *j*.

